# An Immuno-Epidemiological Model of Foot-and-Mouth Disease (FMD) in the African Buffalo Population with Carrier Transmission

**DOI:** 10.1101/2023.09.10.556591

**Authors:** Summer Atkins, Hayriye Gulbudak, J. Shane Welker, Houston Smith

## Abstract

Foot-and-mouth disease (FMD) is a highly contagious disease that spreads among cloven-hoofed animals. Although not deadly, FMD can cause major delays in meat and dairy production. One major concern is that the foot-and-mouth disease virus (FMDV) can persist in African buffalo hosts as natural reservoirs for long periods of time, causing the pathogens to reemerge in susceptible populations. In this paper, we present a novel immuno-epidemiological model of FMD in the African buffalo host populations. Upon infection, the hosts can undergo two phases, namely the acute and the carrier stages. In our model, we divide the infectious population based upon these two stages so that we can dynamically capture the immunological characteristics of both phases of the disease to better understand the carrier’s role in disease transmission. We first define the within-host viral-immune kinetics dependent epidemiological basic reproduction number ***ℛ*** _**0**_ and show that it is a threshold condition for the local stability of the disease-free equilibrium and existence of the endemic equilibrium. We also analytically show that the system always displays forward bifurcation with respect to between-host epidemic parameters. Later, by using a sensitivity analysis (SA) approach developed for multi-scale models, we assess the impact of the acute infection and carrier phase immunological parameters on ***ℛ*** _**0**_. Interestingly, our numerical results show that the within-carrier infected host immune kinetics parameters and the susceptible individual recruitment rates play significant roles in disease persistence, which are consistent with experimental and field studies.

## 1 Introduction

The foot-and-mouth disease virus (FMDV) is a fast-evolving and fast-transmitting pathogen that can infect a plethora of cloven-hoofed species, including domestic livestock and numerous wildlife species (Garabed et al., 2020). Transmission of FMDV can occur through both direct and indirect contact between animals, through fomites, and even through airborne spread under certain atmospheric conditions (Gloster, 1983). Although the foot-and-mouth disease (FMD) is not deadly, outbreaks of FMD can be devastating to agriculture and wildlife economies. The economic impact of FMD is largely due to control measures that are used for curtailing the spread of the disease among livestock populations, *e*.*g*. restricting market access for livestock and livestock products from affected areas, quarantining of infected/exposed animals, culling of infected animals, and even mass culling of contagious premises (Shikumwifa, 2022). FMD remains a serious economic problem in South African regions. The main obstacle in disease eradication is the presence of large numbers of infected African buffalo, as they are able to transfer FMDV to livestock with serotypes South African Territories (SAT) 1, 2, and 3 (Shikumwifa, 2022; Michel and Bengis, 2012). FMD infected buffalo can undergo two stages of the disease: the primary infection stage, which is called the acute phase, and a secondary stage, which is commonly called the carrier phase because they serve as carriers to the disease. In this paper, we construct an immuno-epidemiological model of FMD in the buffalo population which captures the immunological characteristics of the carrier phase to better understand how carrier infection contributes to the spread of the disease.

In the acute phase, the virus is found in both the bloodstream and in the tonsils. Experimental studies show that viral clearance in the bloodstream is obtained within 30 days post infection; however, the viral load may or may not clear in the tonsils during that time frame (Perez-Martin et al., 2022). If the pathogen load in the tonsils does clear within 30 days, then the individual is considered to be recovered and only undergoes the acute stage of the disease. Otherwise, the individual enters the secondary stage of the disease and is classified as being a carrier. The duration of the carrier phase can last from months to even years (Stenfeldt C, 2020; Shikumwifa, 2022).

Understanding the persistence of FMDV in host populations and how the role of carrier transmission contributes to disease persistence is ultimately a multi-scale problem in which both immunological and epidemiological scales need to be investigated (Stenfeldt C, 2020; Garabed et al., 2020; Jolles et al., 2021). In prior works, deterministic models of FMD have been used for understanding the spread of FMD (Pech and McIlroy, 1990; Chowell et al., 2006), ways of controlling the spread of FMD (Gashirai et al., 2020; Shikumwifa, 2022), and the persistence of FMDV among livestock and reservoir populations (Hayer et al., 2018; McLachlan et al., 2019). With regard to analyzing FMD at the immunological scale, there have been some works (Howey et al., 2009, 2012; Macdonald et al., 2022) in which systems of ordinary differential equations (ODEs) were constructed to model the viral and immunological dynamics of FMDV. Howey et al. (2009) constructed a within-host model to capture the viral dynamics of needle-infected pigs and found a trend in which the higher the initial dose the earlier the development of viral detection. In another work, Howey et al. (2012) constructed a within-host model consisting of eight compartments to investigate viral dynamics of FMDV in cattle and found that the within-host model parameters fitted varied from host to host. They assert that a better understanding of the within-host dynamics also provides insights into the dynamics of infectiousness and the transmission of the virus to new hosts. In a prior work (Macdonald et al., 2022), we constructed a within-host model to capture the viral and immunological dynamics of FMDV in African buffalo during the acute phase. The model was fitted to time series data corresponding to buffalo who were infected with one of three strains (SAT1, SAT2, and SAT3) of FMDV. In analyzing this model via across-scale linkages, we find that the viral replication rate in the host highly correlates with the transmission rate on the population scale and that the adaptive immune response activation rate highly correlates with the acute-infection period, making both within-host viral growth rate and the adaptive immune response activation rate crucial parameters in disease dynamics. We find this work as being a stepping stone to multi-scale modeling of FMD in which the within-host dynamics of FMDV can be rigorously linked to epidemiological scales.

To the best of our knowledge, there have not been any multiscale modeling approaches applied to FMD that link the immunological dynamics to the population dynamics. The majority of the literature associated with modeling of FMD focuses on spatial scales of interest (Bradhurst et al., 2015; Ferguson et al., 2001; Hayama et al., 2013; Keeling et al., 2001; Tildesley et al., 2006; Michael J. Tildesley and Woolhouse, 2009). These studies sought to parameterize models that predict disease spread at farm to regional and national scales in order to best evaluate interventions used for limiting transmission and the risk of major epidemics (note the majority of these models are stochastic). However, there have been works in which carrier transmission is investigated (Shikumwifa, 2022; Thomson et al., 1992; Juleff et al., 2008; Maree et al., 2016; Jolles et al., 2021; McLachlan et al., 2019).

In this paper, we present an immuno-epidemiological model of FMD in African buffalo (in particular serotype SAT1) to investigate how the within-host viral and immune kinetics affect the spread of the disease. In particular, we investigate the role of carrier transmission in the context of FMDV disease ecology. We couple an age-structured epidemiological system (where infection age corresponds to time-since-host infection) with two within-host immunological models representing the acute and carrier phases. We perform mathematical analysis on both the within-host model of carrier-infected buffalo and the age-structured PDE epidemic model. In analyzing the within-host model, we define an immunological basic reproduction number, *ℛ* _0_, that also serves as a threshold condition for the local stability of a pathogen-free equilibrium. Additionally, we present existence and stability conditions for two pathogen persisting equilibria. In investigating the between-host model, we define the within-host viral-immune kinetics dependent epidemiological basic reproduction number *ℛ* _0_ and show that it is a threshold condition for the local stability of the disease-free equilibrium and existence of the endemic equilibrium. Additionally, we analytically show that the system always displays forward bifurcation with respect to between-host epidemic parameters. Numerical results suggest that the presence of carrier infected individuals and the birth of new susceptible individuals are the driving influences of disease persistence, which aligns with what was observed in Jolles et al. (2021). Lastly, we use a sensitivity analysis approach (as developed in Gulbudak et al. (2022)), to investigate the impact of the within-host viral and immune kinetics (during both stages of the disease) on the disease outcomes. Interestingly, numerical results suggest that the adaptive immune response activation rate during the acute phase has the largest impact on the epidemiological basic reproduction number, *ℛ* _0_.

The paper is organized as follows. In Section 2, we first present two immunological models of FMDV, representing the virus-immune response dynamics during the acute and carrier phases. We analyze the within-carrier host immune model. We then present the age-since-infection structured epidemiological between-host model. In Section 3, we provide the threshold and bifurcation analysis of the multiscale model. In Section 4, we present numerical simulations of the model and discuss the epidemiological implications. In Section 5, we use a sensitivity analysis approach to investigate the impact of the within-host parameters (in particular carrier transmission parameters) on the epidemiological quantities. In the last section, we summarize our results and give the conclusion.

## 2 An immuno-epidemiological model of FMD with acute and carrier phases

In this section, we present an immuno-epidemiological model that captures the disease spread of foot-and-mouth disease among African buffalo on two scales (namely the immunological and epidemiological). We first discuss our within-host model, which is initially presented in Macdonald et al. (2022), modeling the virus and immune dynamics of the FMDV infected buffalo during the acute phase. Next, we present the within-host model for carrier-infected buffalo and formulate an age-since-infection structured PDE epidemic model. Lastly, we link these within-host models to a population-scale epidemiological PDE model.

### 2.1 Immunological model of an acute-infected individual

Here, we introduce a within-host viral immune kinetics model describing the acute infection stage of infected hosts (formulated first in our prior work Macdonald et al. (2022)). The model considers the interactions between the innate immune response, *I*_*A*_(*τ*), the virus in the bloodstream, *P* (*τ*), and the adaptive immune response *A*(*τ*), where *τ* refers to the time since infection within a host (in days). The model is given in (1) with initial conditions (densities) *I*_*A*_(*τ*_0_) = *I*_*A*0_, *P* (*τ*_0_) = *P*_0_, and *A*(*τ*_0_) = *A*_0_. The innate response is assumed to clear the pathogen at a rate *θ*. We assume that the adaptive response, which is mediated by neutralizing antibodies produced by the host’s B cells, clears the pathogen at rate *ξ*. We assume that the adaptive immune response is activated based upon two processes. One way is independent of the innate response which is represented by *bA*(*τ*)*P* (*τ*) with activation rate *b*. While the other is due to the alarm signals generated by the innate response, which is represented by *aI*(*τ*)*P* (*τ*)*/*(1 + *I*(*τ*)), with activation rate *a* and with half saturating constant scaled to 1. With regard to the pathogen, logistic growth is considered with a replication rate of *r*_*A*_ and a within-host viral carrying capacity in the absence of adaptive immune response, *K*_*A*_. Innate response effectors are always present at a maintenance level Ω*/μ*, where Ω is the maintenance level and *μ* is the immune response decay rate. When the virus invades, the innate response is activated and we represent the activation as a saturating function of virus density with a maximum rate of *ψ* and a half saturation constant *ω*. The within-host acute infection model is as follows:

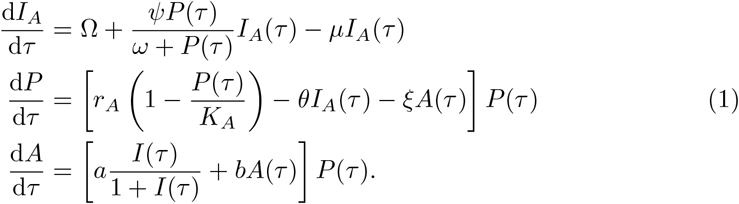

A schematic diagram of the above model is presented on the left-hand side of Figure 1. A detailed analysis of the immunological model is presented in Macdonald et al. (2022). In particular, the following result is established:

**Fig. 1:**
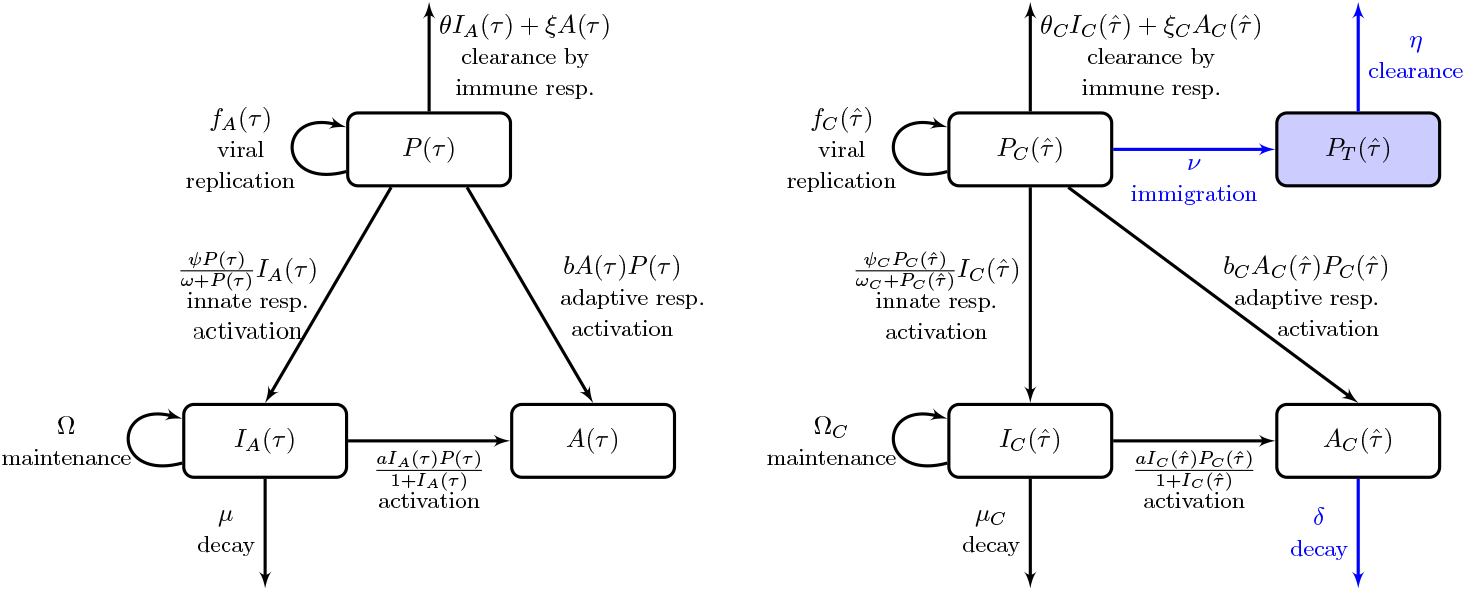
Flowcharts of the within-host dynamics of the acute infectious individual (left) and the carrier infected individual (right), with *f*_*A*_(τ) = *r*_*A*_*P*(τ)[1 − *P*(τ)/*K*_*A*_] and 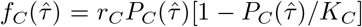. Note that only a proportion of acute-infectedindividuals undergo a carrier phase.

#### Theorem 2.1

(Macdonald et al. (2022)). *If I*_*A*0_ *>* 0 *or A*_0_ *>* 0, *then the pathogen (within the host) eventually clears, i*.*e*.,

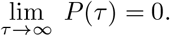

*Subsequently, the innate immune response antibodies decay and the adaptive immune response antibodies increase to a steady state solution, respectively, i*.*e*.,

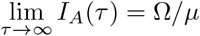

*and*

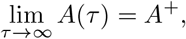

*where A*^+^ *>* 0 *depends on the initial condition; i*.*e. A*^+^ = *z*(*P*_0_, *I*_*A*0_, *A*_0_).

Parameter and state variable descriptions of system (1) are given in Table 3 and Table 4. The acute infection model parameters are estimated in our prior work Macdonald et al. (2022).

**Table 1:**
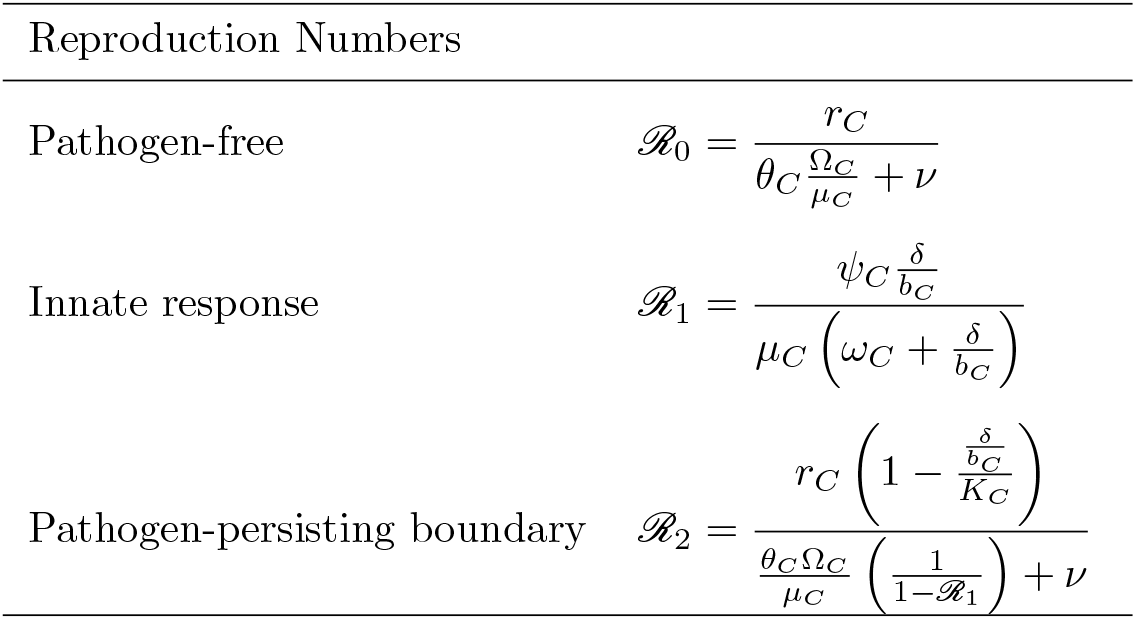
Threshold quantities for the within-carrier system.

**Table 2:**
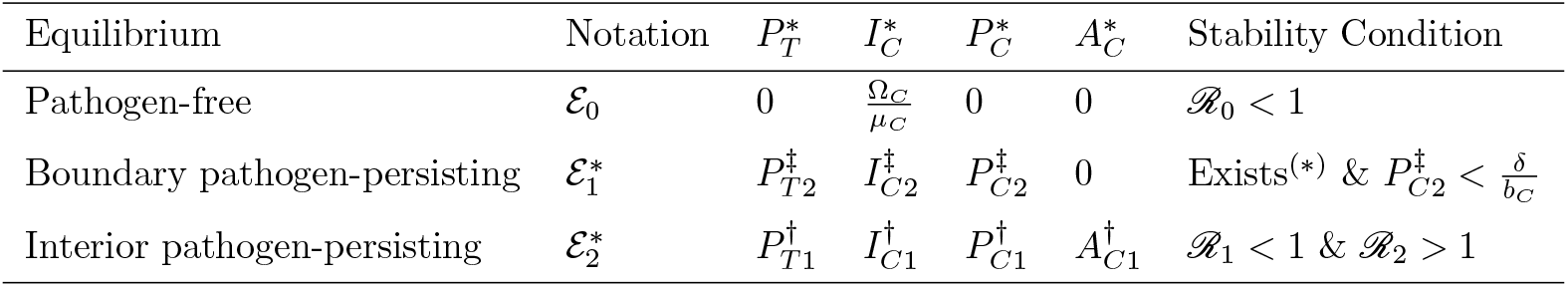
Equilibria and local stability for the within-carrier system.

**Table 3:**
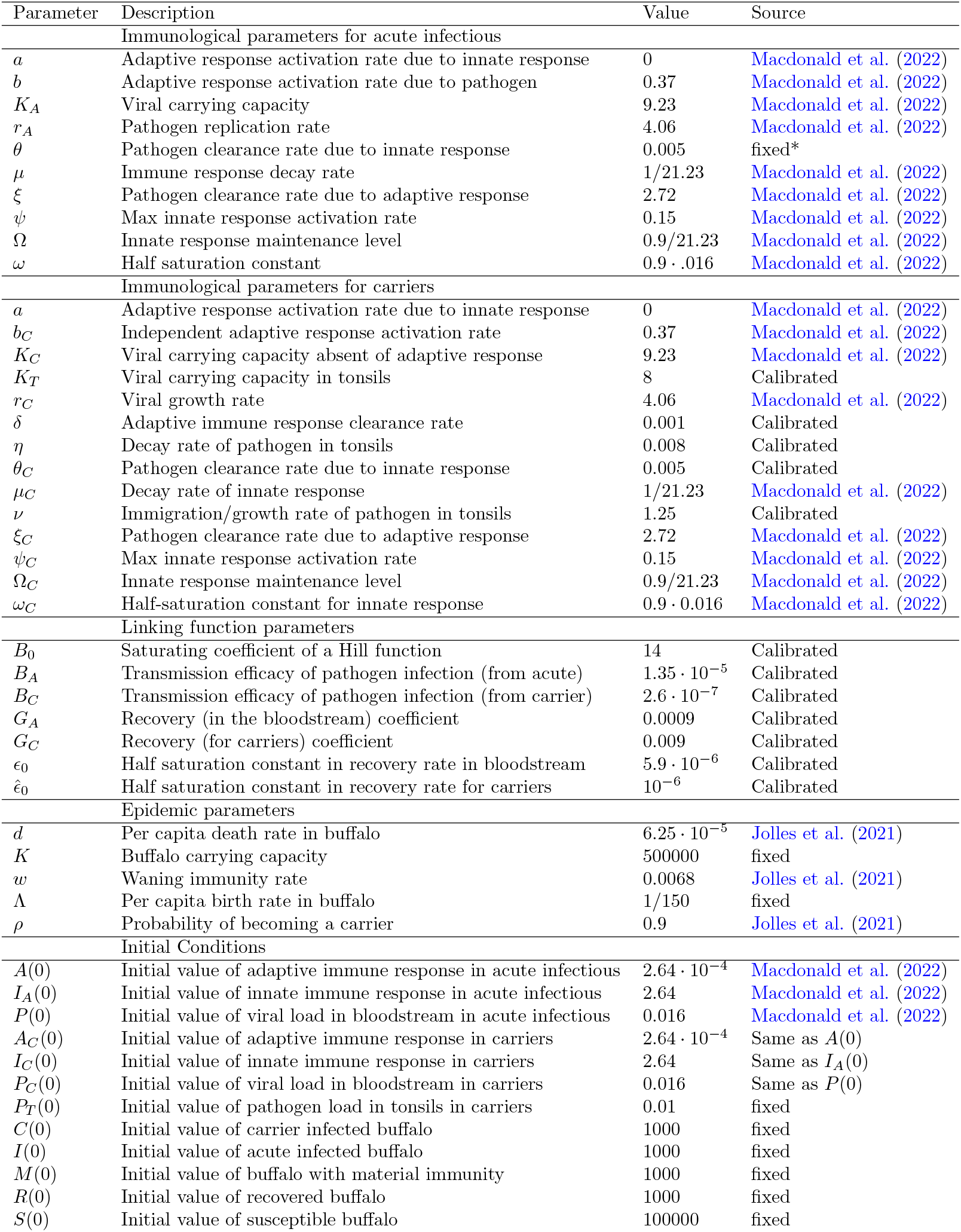
List of parameters along with their descriptions, values, and sources. *-in Macdonald et al. (2022), θ was fixed as being 0, but we fix θ as being the calibrated value for θ_C_ so that *I*_*A*_ isn’t decoupled from system (1).

| Parameter | Description | Value | Source |
| --- | --- | --- | --- |
| Immunological parameters for acute infectious |  |  |  |
| $a$ | Adaptive response activation rate due to innate response | 0 | Macdonald et al. (2022) |
| $b$ | Adaptive response activation rate due to pathogen | 0.37 | Macdonald et al. (2022) |
| $K_A$ | Viral carrying capacity | 9.23 | Macdonald et al. (2022) |
| $r_A$ | Pathogen replication rate | 4.06 | Macdonald et al. (2022) |
| $\theta$ | Pathogen clearance rate due to innate response | 0.005 | fixed* |
| $\mu$ | Immune response decay rate | 1/21.23 | Macdonald et al. (2022) |
| $\xi$ | Pathogen clearance rate due to adaptive response | 2.72 | Macdonald et al. (2022) |
| $\psi$ | Max innate response activation rate | 0.15 | Macdonald et al. (2022) |
| $\Omega$ | Innate response maintenance level | 0.9/21.23 | Macdonald et al. (2022) |
| $\omega$ | Half saturation constant | 0.9 · .016 | Macdonald et al. (2022) |
| Immunological parameters for carriers |  |  |  |
| $a$ | Adaptive response activation rate due to innate response | 0 | Macdonald et al. (2022) |
| $b_C$ | Independent adaptive response activation rate | 0.37 | Macdonald et al. (2022) |
| $K_C$ | Viral carrying capacity absent of adaptive response | 9.23 | Macdonald et al. (2022) |
| $K_T$ | Viral carrying capacity in tonsils | 8 | Calibrated |
| $r_C$ | Viral growth rate | 4.06 | Macdonald et al. (2022) |
| $\delta$ | Adaptive immune response clearance rate | 0.001 | Calibrated |
| $\eta$ | Decay rate of pathogen in tonsils | 0.008 | Calibrated |
| $\theta_C$ | Pathogen clearance rate due to innate response | 0.005 | Calibrated |
| $\mu_C$ | Decay rate of innate response | 1/21.23 | Macdonald et al. (2022) |
| $\nu$ | Immigration/growth rate of pathogen in tonsils | 1.25 | Calibrated |
| $\xi_C$ | Pathogen clearance rate due to adaptive response | 2.72 | Macdonald et al. (2022) |
| $\psi_C$ | Max innate response activation rate | 0.15 | Macdonald et al. (2022) |
| $\Omega_C$ | Innate response maintenance level | 0.9/21.23 | Macdonald et al. (2022) |
| $\omega_C$ | Half-saturation constant for innate response | 0.9 · 0.016 | Macdonald et al. (2022) |
| Linking function parameters |  |  |  |
| $B_0$ | Saturating coefficient of a Hill function | 14 | Calibrated |
| $B_A$ | Transmission efficacy of pathogen infection (from acute) | $1.35 \cdot 10^{-5}$ | Calibrated |
| $B_C$ | Transmission efficacy of pathogen infection (from carrier) | $2.6 \cdot 10^{-7}$ | Calibrated |
| $G_A$ | Recovery (in the bloodstream) coefficient | 0.0009 | Calibrated |
| $G_C$ | Recovery (for carriers) coefficient | 0.009 | Calibrated |
| $\epsilon_0$ | Half saturation constant in recovery rate in bloodstream | $5.9 \cdot 10^{-6}$ | Calibrated |
| $\hat{\epsilon}_0$ | Half saturation constant in recovery rate for carriers | $10^{-6}$ | Calibrated |
| Epidemic parameters |  |  |  |
| $d$ | Per capita death rate in buffalo | $6.25 \cdot 10^{-5}$ | Jolles et al. (2021) |
| $K$ | Buffalo carrying capacity | 500000 | fixed |
| $w$ | Waning immunity rate | 0.0068 | Jolles et al. (2021) |
| $\Lambda$ | Per capita birth rate in buffalo | 1/150 | fixed |
| $\rho$ | Probability of becoming a carrier | 0.9 | Jolles et al. (2021) |
| Initial Conditions |  |  |  |
| $A(0)$ | Initial value of adaptive immune response in acute infectious | $2.64 \cdot 10^{-4}$ | Macdonald et al. (2022) |
| $I_A(0)$ | Initial value of innate immune response in acute infectious | 2.64 | Macdonald et al. (2022) |
| $P(0)$ | Initial value of viral load in bloodstream in acute infectious | 0.016 | Macdonald et al. (2022) |
| $A_C(0)$ | Initial value of adaptive immune response in carriers | $2.64 \cdot 10^{-4}$ | Same as $A(0)$ |
| $I_C(0)$ | Initial value of innate immune response in carriers | 2.64 | Same as $I_A(0)$ |
| $P_C(0)$ | Initial value of viral load in bloodstream in carriers | 0.016 | Same as $P(0)$ |
| $P_T(0)$ | Initial value of pathogen load in tonsils in carriers | 0.01 | fixed |
| $C(0)$ | Initial value of carrier infected buffalo | 1000 | fixed |
| $I(0)$ | Initial value of acute infected buffalo | 1000 | fixed |
| $M(0)$ | Initial value of buffalo with material immunity | 1000 | fixed |
| $R(0)$ | Initial value of recovered buffalo | 1000 | fixed |
| $S(0)$ | Initial value of susceptible buffalo | 100000 | fixed |

**Table 4:** Description of state variables and time variables.

| Variable | Description |
| --- | --- |
| $\tau$ | Infection age: Time-since-host-infection for acute-infected individuals |
| $P(\tau)$ | Pathogen load in bloodstream of an acute infectious host with infection age $\tau$ |
| $I_A(\tau)$ | Innate immune response of an acute infectious host with infection age $\tau$ |
| $A(\tau)$ | Adaptive immune response of an acute infectious host with infection age $\tau$ |
| $\hat{\tau}$ | Carrier age: Time since carrier-infected host became a carrier |
| $P_T(\hat{\tau})$ | Pathogen load in the tonsils of a carrier host with carrier age $\hat{\tau}$ |
| $P_C(\hat{\tau})$ | Pathogen load in the bloodstream of a carrier host with carrier age $\hat{\tau}$ |
| $I_C(\hat{\tau})$ | Innate immune response of a carrier host with carrier age $\hat{\tau}$ |
| $A_C(\hat{\tau})$ | Adaptive immune response of the carrier host with carrier age $\hat{\tau}$ |
| $t$ | Chronological time or time of the epidemic |
| $M(t)$ | The number of individuals with maternal immunity at time $t$ |
| $S(t)$ | The number of susceptible individuals at time $t$ |
| $i(t, \tau)$ | The density of acute infectious individuals with infection age $\tau$ at time $t$ |
| $R(t)$ | The number of recovered individuals at time $t$ |
| $c(t, \hat{\tau})$ | The density of the carrier individuals with carrier age $\hat{\tau}$ at time $t$ |

Figure 2 displays a time-dependent solution of the within-acute model (1) (with parameter value settings given in Table 3) which was numerically obtained through MATLAB’s ODE solver ode45. Notice that the pathogen in the bloodstream peaks around day 3. Also the time *τ* in which *P*_*A*_(*τ*) *<* 10^*−*4^ (the lowest viral density threshold for assuming pathogen clearance) is approximately 7.7 days. This is reasonably close to estimates obtained from Jolles et al. (2021) which approximated the acute infection period at 5.7 days. Once the pathogen has been cleared from the blood-stream, the individual either recovers or progresses to the carrier infection stage. The latter case is due to the pathogen persisting in the tonsils.

**Fig. 2:**
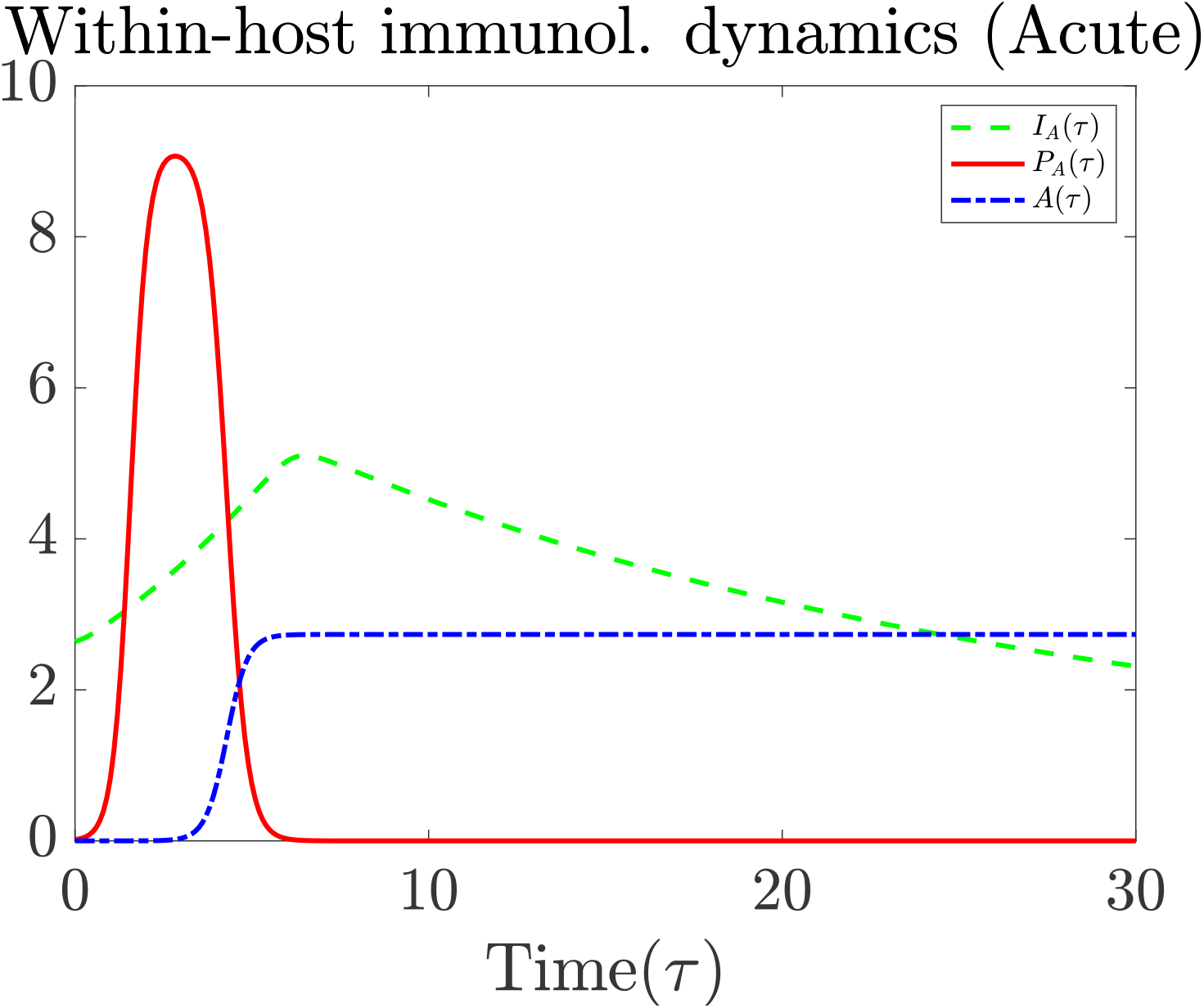
Within-host pathogen-immune response antibody dynamics of an infectious individual during the acute phase. Parameter values used are given in Table 3.

### 2.2 Immunological model of a carrier-infected individual

Here, we formulate an immunological model, describing the within-carrier host virus-immune response dynamics. Similar to the within-host model for the acutely infectious individuals, the model variables are the innate immune response, 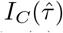, the viral load in the bloodstream, 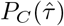, and the adaptive immune response, 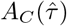, with the independent time variable, 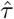, referring to the time passed since infection. Additionally, the within-carrier model has a compartment describing the pathogen load in the tonsils, 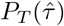. Regarding FMD infection, the pathogen reaches the tonsils by way of dendritic cell transportation, and an FMD-infected buffalo becomes a carrier when the pathogen clears in the bloodstream but remains in the tonsils for an extended period of time.

For FMD, unlike many other pathogens, dendritic cells can pick up complete copies of the virus instead of only viral fragments. Consequently, we assume the pathogen enters the tonsils at rate 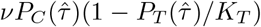. The parameter *K*_*T*_ corresponds to a viral carrying capacity in the tonsils. We also assume the pathogen in the tonsils exits at rate *η*, at which point it may be transported to mucosal cells allowing for the possibility of carrier transmission. Since the carrier phase is significantly longer than the acute phase, we add a decay rate for the adaptive immune response, *δ*, to the model. We also assume that the adaptive response is activated through two processes: (1) a process that is independent of the innate immune response at a rate *b*_*C*_, and (2) a process that depends on the innate immune response. The activation rate that depends on the innate immune response is represented by *aI*_*C*_(*τ*)*/*(1 + *I*_*C*_(*τ*)), but like the within-host acute model, we set *a* = 0. We assume that innate immune response effectors are always present at a maintenance level Ω_*C*_ in the absence of the virus. The innate immune response is activated at a maximum rate *ψ*_*C*_ with half-saturation constant, *ω*_*C*_, in terms of the viral load and decays at a rate *μ*_*C*_.

The model is given in (2) with initial conditions (densities) being 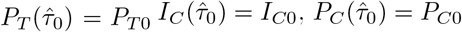, and 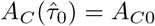.

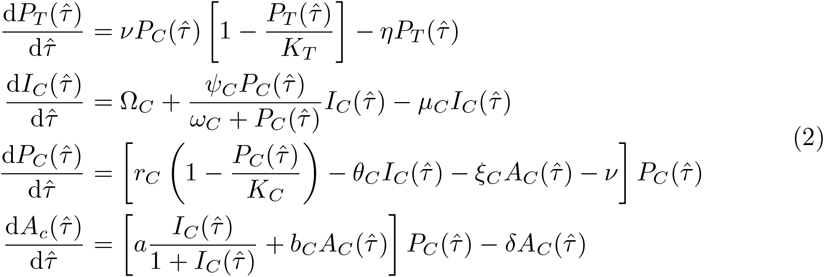

A schematic diagram of the above within-host model for the carrier stage is provided on the right-hand side of Figure 1.

Similar to the within-host acute model (1), logistic growth is assumed for the pathogen in the bloodstream with a replication rate *r*_*C*_ and viral carrying capacity *K*_*C*_ in the absence of the adaptive immune response. Additionally, we assume that the innate response clears the pathogen at a rate *θ*_*C*_, and the adaptive immune response clears the pathogen at a rate *ξ*_*C*_ due to neutralizing antibodies produced by the host’s B cells. In regards to the parameter settings, we set the value for Ω_*C*_, *ψ*_*C*_, *ω*_*C*_, *μ*_*C*_, *r*_*C*_, *K*_*C*_, *θ*_*C*_, *ξ*_*C*_, *b*_*C*_ equal to Ω, *ψ, ω, μ, r*_*A*_, *K, θ, ξ*, and *b* (respectively), the parameter values from acute model. The data that we use to calibrate the remaining parameters come from experiments generated in Perez-Martin et al. (2022).

A detailed analysis of the immunological model (2) is presented below and proofs are provided in Appendix A. It is worth noting that system (2) (with *a* = 0) possesses a pathogen-free equilibrium (PFE) and up to two pathogen-persisting equilibria. Before presenting the analytical results, we define the immunological basic reproduction number

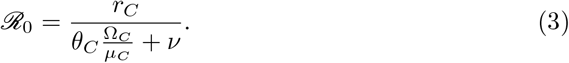

The immunological reproduction number, *ℛ*_0_, gives the number of secondary viral particles that one viral particle will produce in the bloodstream (before the adaptive immune response is mounted). Next we establish the following result:

#### Theorem 2.1.

*The pathogen-free equilibrium*

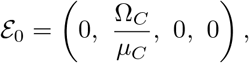

*which always exists, is locally asymptotically stable when*

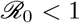

*and unstable when*

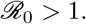

*Proof*. See the proof in Appendix A.1.

In the absence of an adaptive immune response, the system (2) also has a pathogen-persisting boundary equilibrium

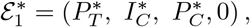

where

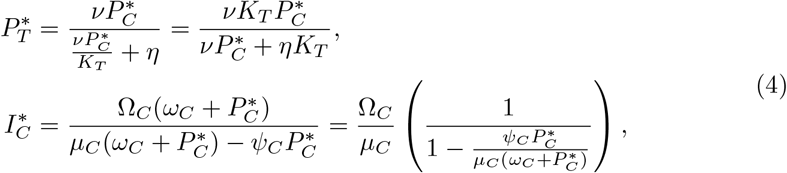

and 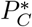 is the positive root to the following quadratic equation:

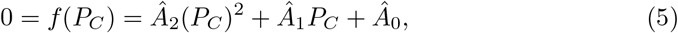

where

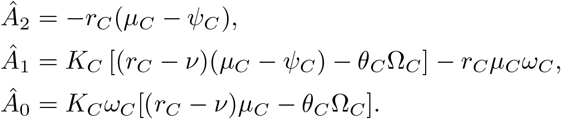

Note that we can rewrite the coefficients for *Â* _1_ and *Â* _0_ as follows:

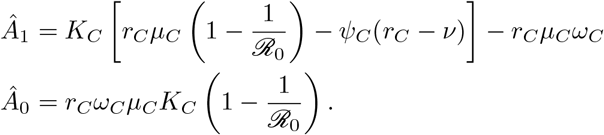

#### Theorem 2.2.

*If one of the two following conditions below holds:*

1. *ℛ*_0_ *>* 1 *and μ*_*C*_ *> ψ*_*C*_.
2. *ℛ*_0_ *>* 1, *μ*_*C*_ *< ψ*_*C*_, *and the discriminant of the quadratic polynomial f* (*P*_*C*_), 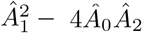, *is positive*,

*then there exists a unique pathogen-persisting boundary equilibrium* 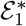. *Moreover, if* 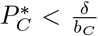, *then the pathogen persisting boundary equilibrium is locally asymptotically stable*.

*Proof*. See the proof in Appendix A.2.

In addition, the system (2) has a pathogen-persisting interior equilibrium

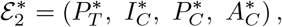

where

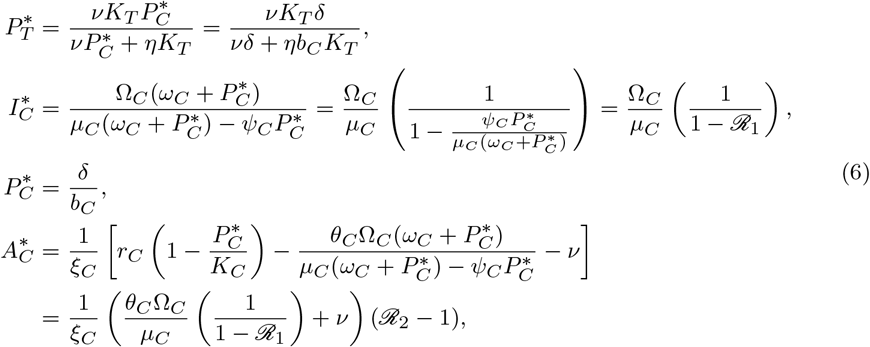

where

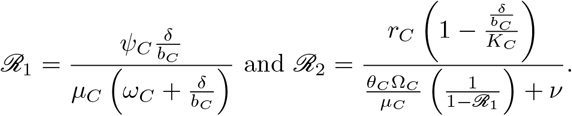

We establish the following analytical result:

#### Theorem 2.3.

*The pathogen-persisting interior equilibrium*, 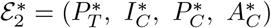, *exists and is locally asymptotically stable if*

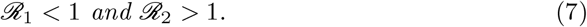

*Proof*. See the proof in Appendix A.3.□

It is worth mentioning that the two inequalities in (7) imply that we are in a parameter setting in which *ℛ*_0_ *>* 1 (i.e. the pathogen-free equilibrium is unstable).

Figure 3a displays the solution of the system (2) with parameter settings given in Table 3. Note that the darker gray region represents the acute phase of the infected individual while the lighter gray region is the carrier phase. In Figure 3b, we see the same plot of the carrier phase over the first 750 days. We only consider the first 750 days since the data (Perez-Martin et al., 2022) used for parameterization involve juvenile buffalo who are only observed for one to three years. The majority of the parameter values are obtained from Macdonald et al. (2022). However, based upon these parameter values, the solution to the dynamical system converges to the pathogen-persisting interior equilibrium, with 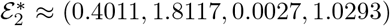. Figure 11 in Appendix D illustrates that the carrier individual eventually experiences multiple instances of viral relapse, where the solution converges to the pathogen-persisting interior equilibrium.

**Fig. 3:**
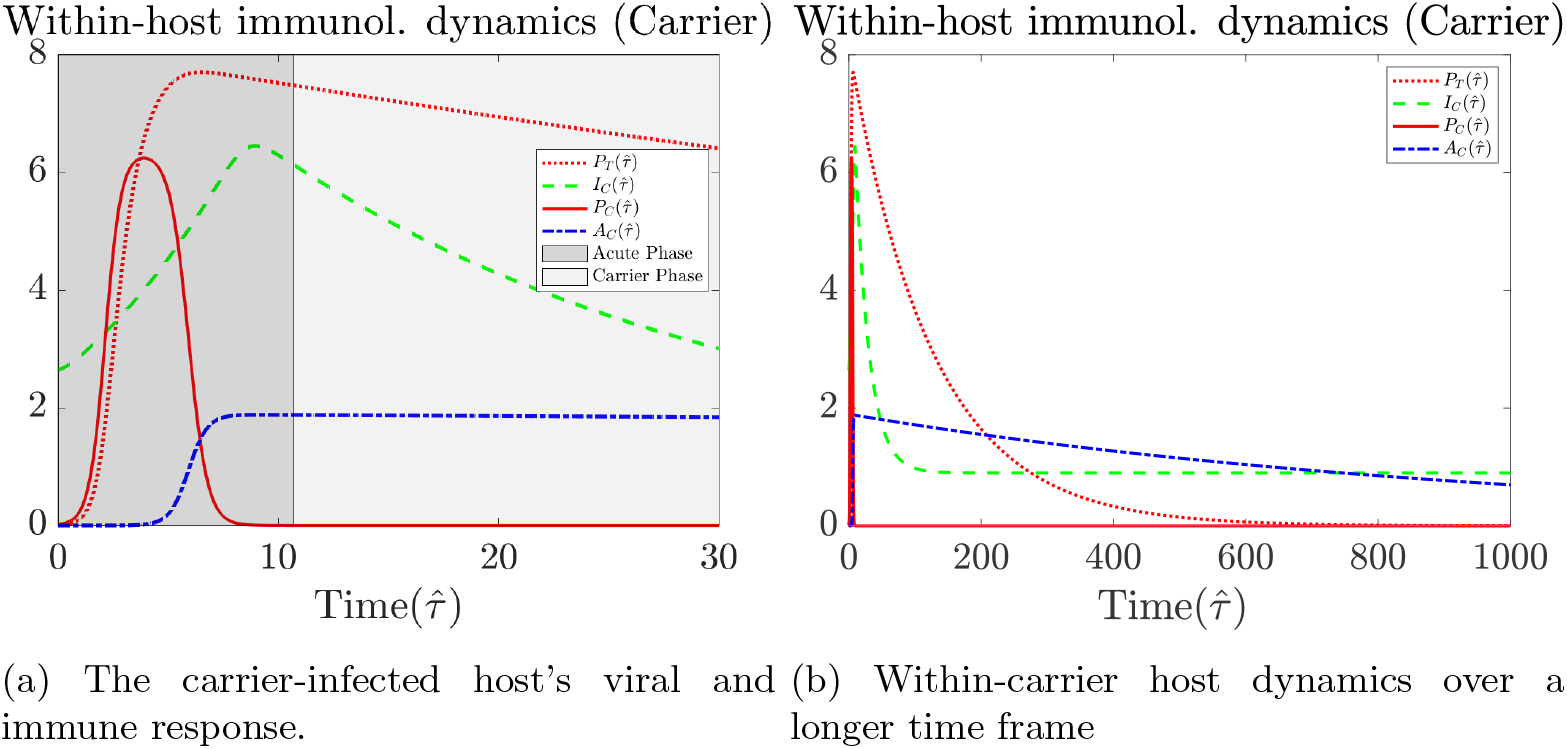
The within-host dynamics of an infected buffalo during the carrier stage. Parameter values used are given in Table 3. Note that carrier-infected hosts experience an acute phase (the dark grey shaded region in subfigure (a)) prior to carrier phase. Subfigure (b) presents the same dynamics but over a longer time frame, illustrating that the pathogen load in the tonsils starts to become negligible around 500 days.

**Fig. 4:**
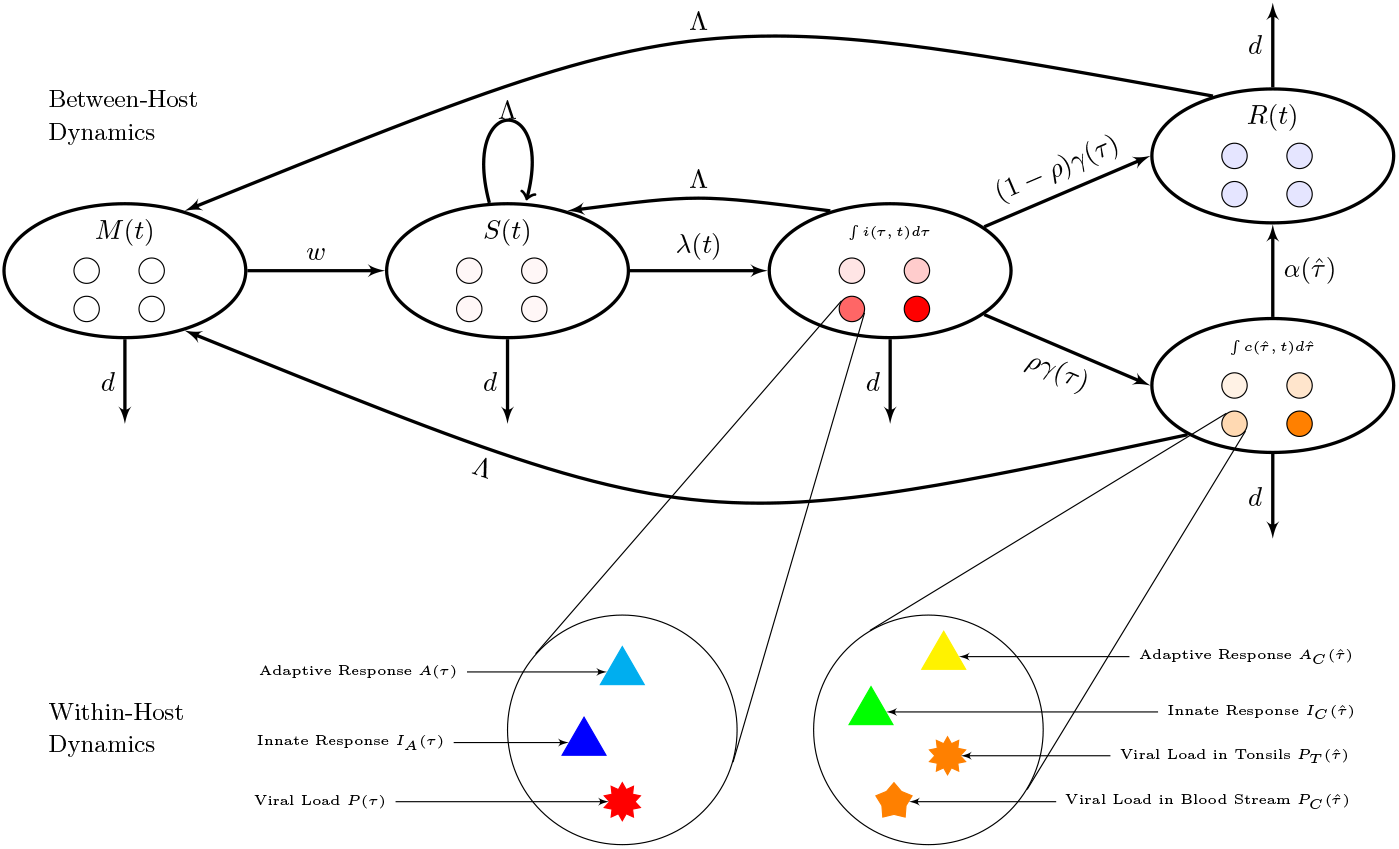
Flowchart of the multi-scale model. Here λ(t) represents the force of infection due to interacting with either an acute infected individual or a carrier infected individual.

### 2.3 The epidemiological model

At the *population scale*, we consider a time-since-infection structured epidemic model with the acute and carrier stages of infection. In the model, the variables *S*(*t*), *I*(*t*), *C*(*t*), *R*(*t*), and *M* (*t*) represent the numbers of *susceptible, acute-infected, carrier-infected, recovered*, and the *maternally immune* host populations, respectively. Let *i*(*τ, t*) represents the acute-infected host density with time-since-host infection *τ* at time *t*. Let 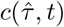 represent the carrier infected-host density, where 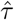 is the time since becoming a carrier. The total number of infected individuals during the acute phase at time *t* is

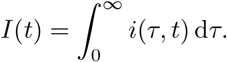

Similarly, the number of infected individuals that are carriers at time *t* is

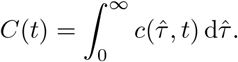

The full model of the between-host system is as follows:

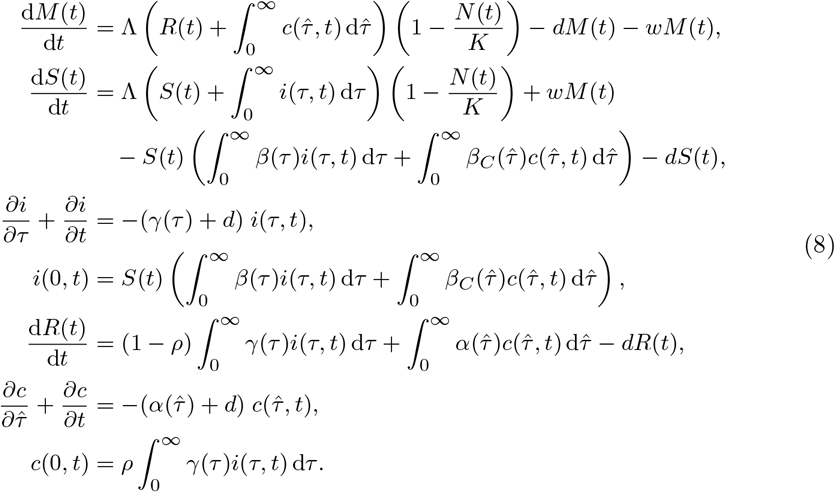

Animals in the recovered (*R*) and carrier (*C*) classes give birth to individuals who enter the *M* class due to obtaining maternal immunity. Animals belonging to other classes give birth to new susceptible individuals. Here, *N* (*t*) represents the total host population. We considered logistic growth rates, where Λ represents the per capita birth rate of buffalo and *K* represents the carrying capacity. Maternally immune individuals eventually exit the class due to their immunity waning at rate *w* and enter the susceptible class. We assume a density-dependent force of infection from infected individuals during the acute phase 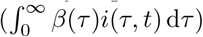 and from carrier-infected individuals during the carrier phase 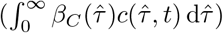 After successful interactions with an acutely infected or carrier infected buffalo, susceptible individuals move to infectious class *i*(*t, τ*) with a rate

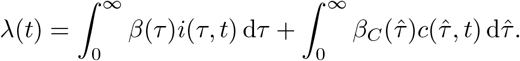

Upon clearing the virus in the bloodstream, if the hosts still has the pathogen present in the tonsils, we assume these individuals become carriers and move to the carrier phase. However, not all acutely infected buffalo transition into the carrier stage. For this model, we assume that the acutely infected hosts exit at a rate *γ*(*τ*), which represents the clearance rate of the pathogen in the bloodstream. We let 0 *< ρ <* 1 be the fraction of the acute infected individuals who become carriers, and (1 *− ρ*) is the fraction of the acute infected individuals who enter the recovered class. The carriers eventually recover at rate 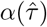. *In vivo* studies, along with a modeling framework that we develop here, allow us to bridge the immunological and population scales by defining functional forms of epidemic parameters as functions of the within-host virus immune response. As suggested by data in Handel and Rohani (2015); Fraser et al. (2014); Gulbudak et al. (2017), to bridge the scales from individual to population, the unit infectiousness and recovery rates can be formulated as functions of immunological variables as follows:

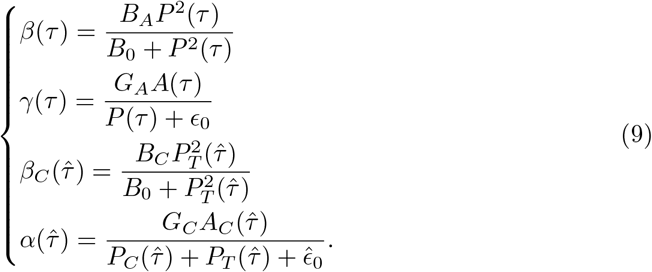

The transmission rate due to interaction with an acutely infected individual, *β*(*τ*), is modeled by a sigmoidal Hill equation of *n* = 2 that depends on the pathogen load in the bloodstream. The transmission rate due to interaction with a carrier infected individual, *β*_*C*_(*τ*), is also modeled by a sigmoidal Hill equation of *n* = 2, but this function depends on the viral load in the tonsils. A Michaelis-Menton equation with *n* = 1 is considered for modeling the rate at which the viral load in the bloodstream is cleared, *γ*(*τ*) (Gilchrist and Sasaki, 2002; Gulbudak and Browne, 2020; Tuncer et al., 2016). The recovery rate for the carriers, *α*(*τ*), has a form similar to *γ*(*τ*) but depends on both the viral load in the bloodstream and the tonsils. The parameters used for these linking functions are described in Table 3.

## 3 Equilibria and the threshold analysis

In this section, we first define the within-host virus-immune kinetics-dependent epidemiological basic reproduction number (10) and show that it is a threshold condition for both the stability of the disease-free equilibrium (DFE) and the existence of the endemic (EE) equilibrium.

The basic reproduction number, *ℛ*_0_, of the epidemiological model (8) is

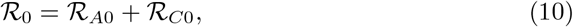

where

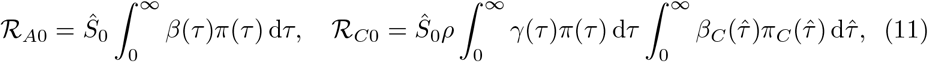

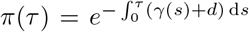 gives the probability of still being in the acute phase *τ* days after becoming infected, and 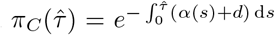 is the probability of still being in the carrier phase 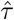days after becoming a carrier.

The first term *ℛ*_*A*0_ accounts for the total number of secondary cases that is produced by one infectious individual (while in the acute phase) in an entirely susceptible population. The second term *ℛ*_*C*0_ accounts for the total number of secondary cases produced by one infected individual (while in the carrier phase) in a population consisting only of susceptible individuals.

### 3.1 Existence and the stability of the DFE

We derive the steady state solutions to the epidemiological model (8) by setting the derivatives with respect to *t* equal to zero:

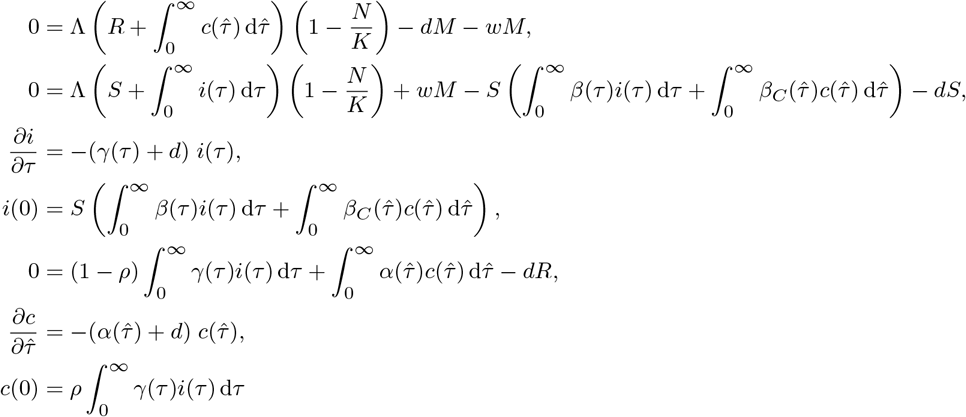

The system (8) has a DFE

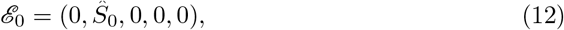

where *Ŝ*_0_ = *K* (1 *− d/*Λ).

Next, we show that ℛ_0_ is a threshold condition for the local stability of the DFE:

#### Theorem 3.1.

*The disease-free equilibrium ℰ*_0_ *(12) is locally asymptotically stable if* ℛ_0_ *<* 1. *If ℛ*_0_ *>* 1, *it is unstable*.

*Proof*. We linearize the system around the DFE, *E*_0_, to determine conditions for stability. We consider solutions near the DFE by setting

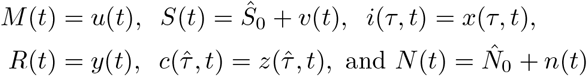

where 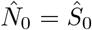. Substituting the perturbed solutions into equation (8), we obtain the following linearized system:

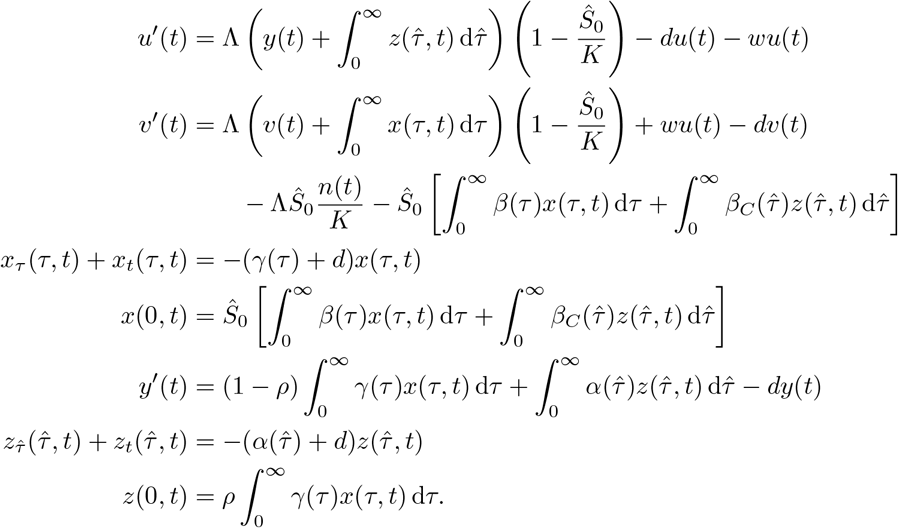

We then seek solutions to the above system that are of the form *u*(*t*) = *ūe*^*λt*^, 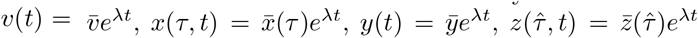, and 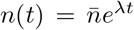 where *λ* is either real or complex. Substituting the time-dependent solutions with the separable ones and canceling *e*^*λt*^, we obtain the following linear eigenvalue problem:

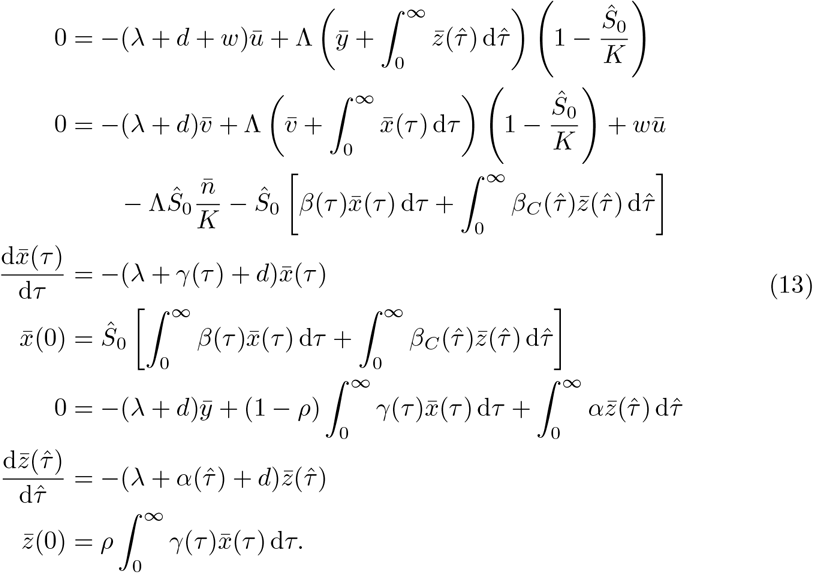

From the fourth equation in (13), we obtain

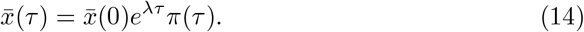

By the sixth equation in (13), we have

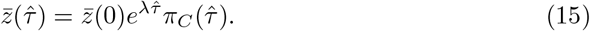

By substituting (14) into the last equation of the system (13), we get a solution for 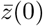 which is a function of 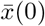:

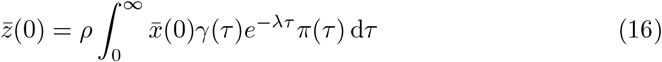

By substituting (14)-(16) into the third equation in (13) and then canceling 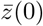, we obtain the characteristic equation *𝒢* (*λ*) = 1, where

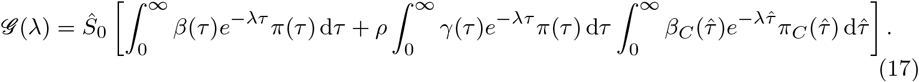

Note that *ℛ* _0_ = *𝒢* (0), where

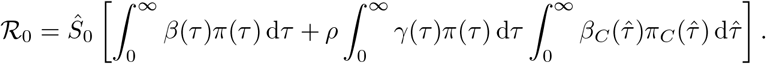

The DFE, *ℰ*_0_, that is given in (12) is locally asymptotically stable if all eigenvalues *λ* have negative real parts, and *ε*_0_ is unstable if there exist a solution *λ* with positive real part.

Let *λ ∈* ℝ and consider the function *𝒢* (*λ*) given in (17). If there is some positive interval for which *β*(*τ*) is strictly positive or if there is some positive function for which both 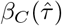 and *γ*(*τ*) are strictly positive, then

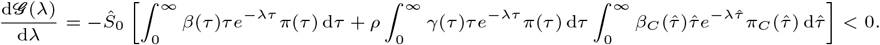

Thus, *𝒢* is a decreasing function of *λ*, and

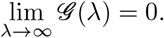

When *ℛ*_0_ = *𝒢* (0) *>* 1, there exists a unique positive real solution to the equation *𝒢* (*λ*) = 1 by Intermediate Value Theorem. Therefore *ℛ*_0_ *>* 1 implies that *ε*_0_ is unstable.

Next, we consider the case where *ℛ*_0_ = *𝒢* (0) *<* 1. Let *λ* = *a* + *bi* be an eigenvalue with nonnegative real part *a*. Then

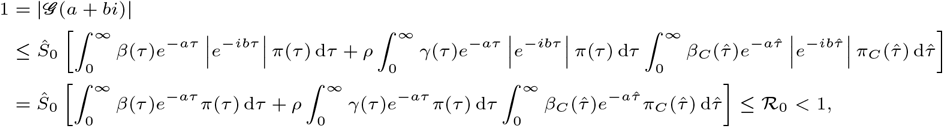

which is a contradiction. Therefore, all eigenvalues have negative real parts when *𝒢*_0_ *<* 1. This completes the proof.

### 3.2 Existence of the endemic equilibrium

Next, we will show the existence of an endemic (positive) equilibrium *ε* ^*\**^ = (*M*^*\**^, *S*^*\**^, *i*^*\**^(*τ*), *R*^*\**^, *c*^*\**^(*τ*)) of the system (8) when *ℛ*_0_ *>* 1. First, we establish the following result:

#### Lemma 3.2.

*Let* 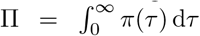, *and* 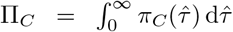 *(where* 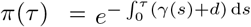*and* 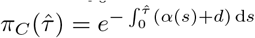).*Define*

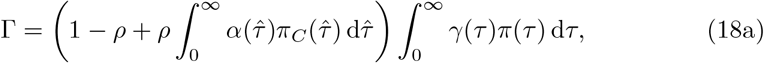

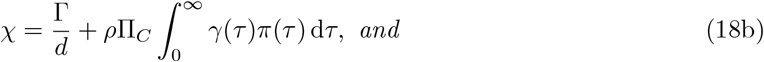

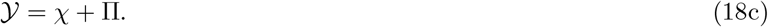

*We can simplify χ and 𝒴 as follows:*

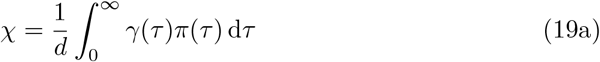

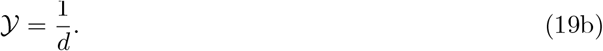

*Proof*. Observe that 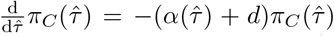 and 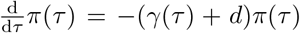. We can use these derivatives to simplify *χ* and *𝒴* (Note: for simplifying χ we also use (18a)).

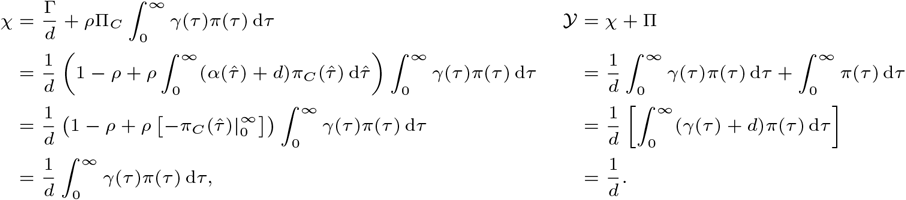

#### Theorem 3.3.

*If ℛ*_0_ *>* 1, *then there exists a unique endemic equilibrium, ε, to system (8) with its components being the following:*

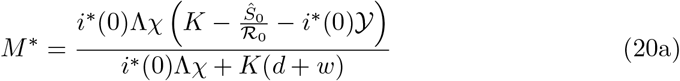

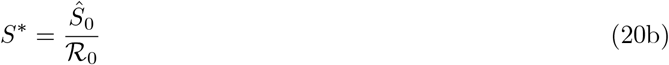

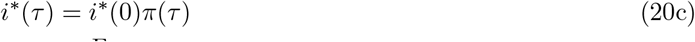

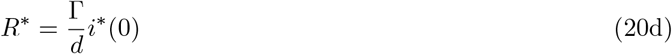

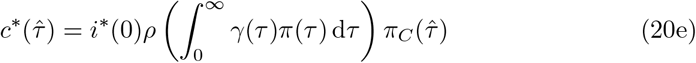

*where* Γ, *χ, and 𝒴 are the numbers defined in Lemma 3*.*2*.

*For the components M*^*\**^, *i*^*\**^(*τ*), *R*^*\**^, *and* 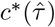, *the component i*^*\**^(0) *must be the root of the following quadratic polynomial*

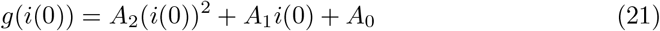

*with coefficients*

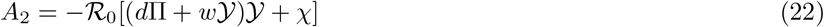

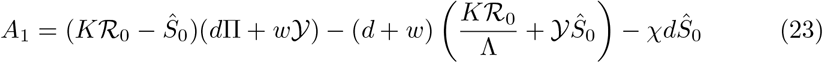

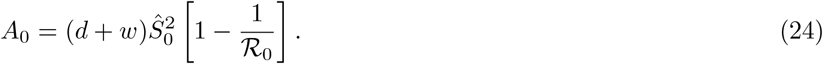

*Proof*. The equilibria of the epidemiological model are obtained by setting the time *t* derivatives of the model to zero:

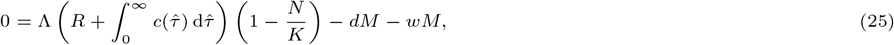

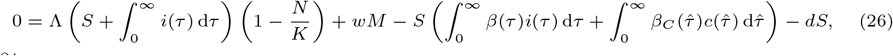

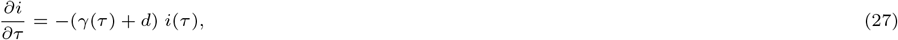

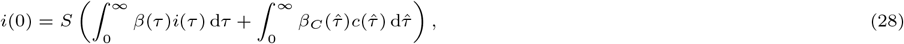

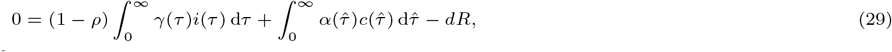

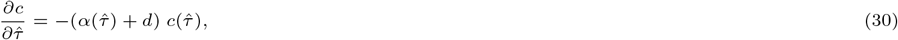

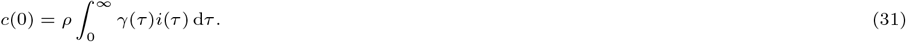

By first solving the differential equation (27), we obtain:

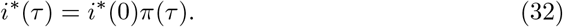

Similarly, we solve the differential equation (30)

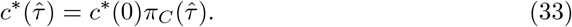

Then substituting (32) into equation (31), we get

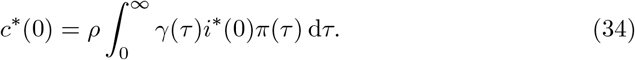

Substituting the expressions for *i*^*\**^(*τ*), 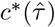, and *c*^*\**^(0) into equation (28) and canceling the *i*^*\**^(0) yields the expression below:

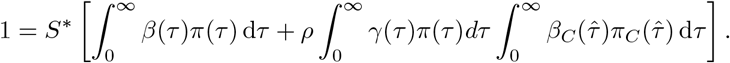

We rearrange terms in the above equation and use (10) to obtain the following:

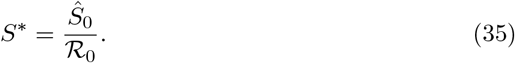

From equation (29), we have

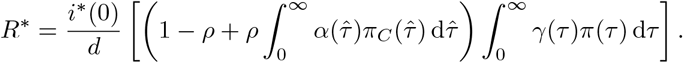

Now, we substitute the expression of Γgiven in (18a) to rewrite *R*^*\**^ as follows:

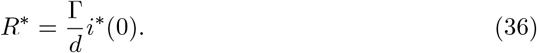

Note that in solving equation (25), we obtain

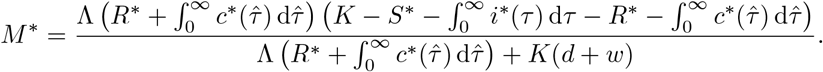

Also note that

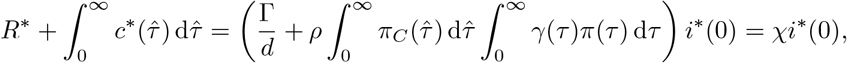

where *χ* is given in (18b). Using equations (32), (35), (18c), and the above equality, we re-express *M*^*\**^ in terms of *i*^*\**^(0)

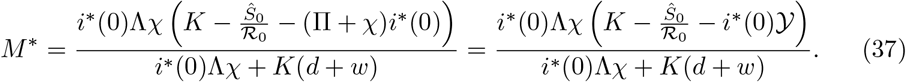

Now, we utilize equation (26) to find *i*^*\**^(0). First, we use (28) to rewrite equation (26)

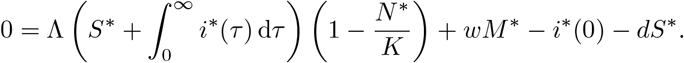

Observe that

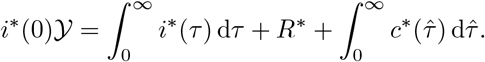

Thus, *N*^*\**^ = *S*^*\**^ + *i*^*\**^(0) *𝒴* + *M*^*\**^. Multiplying both sides of the above equation by *K* and re-expressing *N*^*\**^ yields

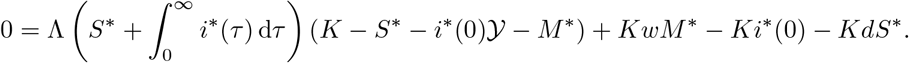

We substitute expressions for *S*^*\**^, *M*^*\**^, and *i*^*\**^(*τ*) into the above equation to obtain the following:

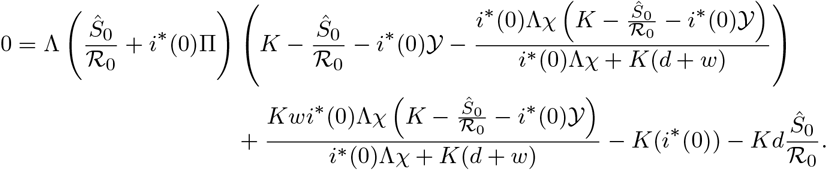

Multiplying both sides of the above equation by *i*^*\**^(0)Λ_*χ*_ + *K*(*d* + *w*) and *ℛ*_0_ and dividing both sides by *K* yields:

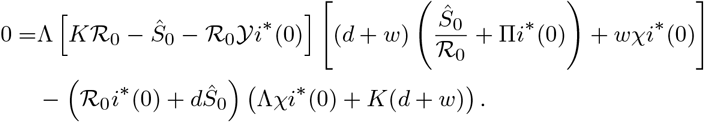

Distributing terms and using *𝒴* = *χ* + Π yields

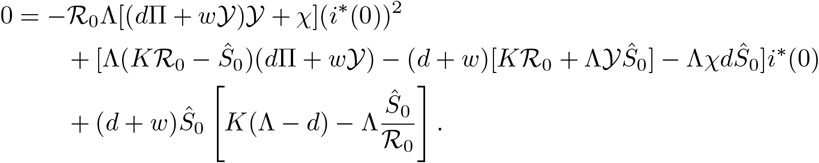

We rewrite the constant term in the above equation by using *Ŝ*_0_ = *K*(Λ *− d*)*/*Λ and divide both sides of the above equation by Λ to obtain

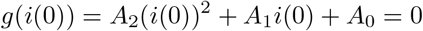

with coefficients

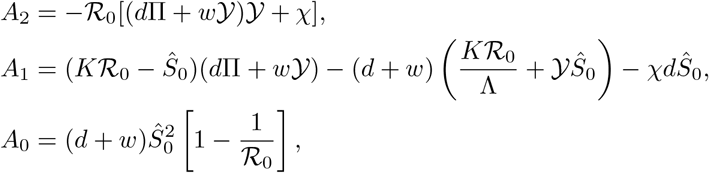

which are the coefficients (22)-(24) of the polynomial (21). Notice that the sign of *A*_2_ is always negative and that the sign of *A*_0_ is positive whenever *ℛ*_0_ *>* 1. Then, by Descartes’ Rule of Signs, *ℛ*_0_ *>* 1 implies that polynomial *g*(*i*(0)) has exactly one positive root, which is of the form

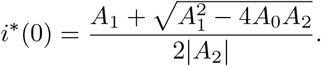

Lastly, we show that the unique positive root satisfies

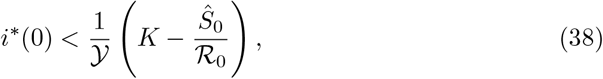

which ensures that *M*^*\**^ is positive. Observe when *ℛ*_0_ *>* 1, the graph of the quadratic polynomial *g* is a parabola that opens down with the *y*-intercept lying above the *x*-axis. Therefore, for verifying inequality (38), it suffices to show that

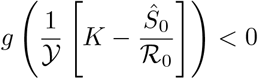

when *ℛ*_0_ *>* 1. We evaluate *g* at (1*/𝒴*)[*K − Ŝ*_0_*/ℛ*_0_], simplify, and rearrange terms to obtain

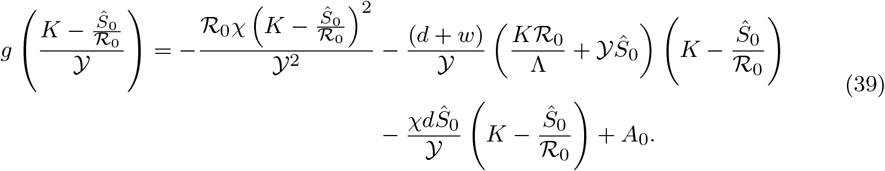

Note that the term (*K − Ŝ*_0_*/ℛ*_0_) *>* 0 since *Ŝ*_0_ *< K* and 1*/ℛ*_0_ *<* 1 when *ℛ*_0_ *>* 1. All terms of the right-hand side of the above equality are negative except the last term, *A*_0_. We combine the second term with the last term (but replace *A*_0_ with (24)) of the right-hand side of (39) with the last term (substituting to form the following expression:

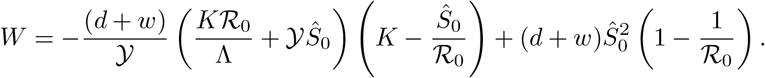

We show that the sign of *W* is negative to verify that the right side of (39) is negative. We use *Ŝ* _0_ = *K*(1 *− d/*Λ) to simplify the expression *W* to get

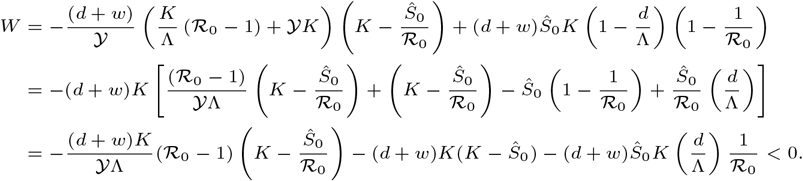

This completes our proof.

The presence of the quadratic equation (21) causes one to consider the plausibility of backward bifurcation. In our case, we have shown uniqueness of an endemic equilibrium when *ℛ* _0_ *>* 1. However, the system may have up to two interior endemic equilibria when *ℛ* _0_ *<* 1, in which case the system may display backward bifurcation with important biological implications (Barfield et al., 2018; Gulbudak et al., 2021; Gulbudak and Martcheva, 2013; Welker and Marcheva, 2020). Thus, we explore the bifurcation dynamics of the system and show that it can present forward bifurcation for some epidemiological parameters. To investigate bifurcation, we consider three parameters, *p ∈ {ρ, K*, Λ*}*, that appear explicitly in *ℛ* _0_. We view *p* as being a potential bifurcation parameter. In order to investigate how the dynamics change with respect to changes in *p*, we consider thresholds as functions of *p* and solve *ℛ*_0_(*p*) = 1 and denote the critical value as being *p*_crit_. In particular, we have

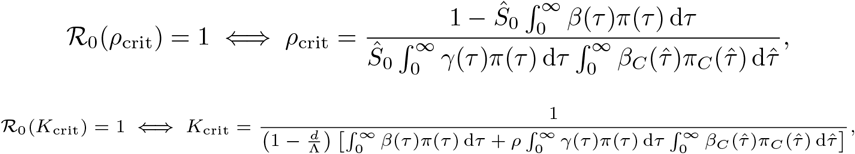

and

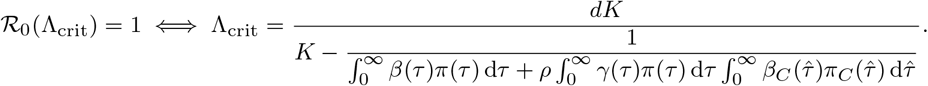

In the latter case, the value of Λ_crit_ is positive since the number

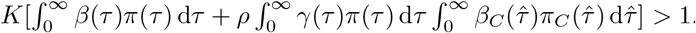

We use the Implicit Function Theorem to show that only forward bifurcation occurs for these parameters at the critical parameter values.

#### Theorem 3.4.

*The system (8) has forward bifurcation at p* = *p*_*crit*_, *I*^*\**^ = 0 *(where p*_*crit*_ *is the critical parameter value that gives ℛ* _0_(*p*_*crit*_) = 1 *and p ∈ {ρ, K*, Λ*})*.

*Proof*. Let *p* be our bifurcation parameter and let *p*_crit_ our critical value for which *ℛ*_0_ (*p*_crit_) = 1. Now let 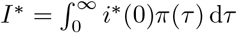 . We look at the sign of

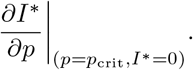

The system has backward bifurcation at *p* = *p*_crit_ if and only if

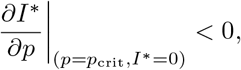

and the system has forward bifurcation at *p* = *p*_crit_ if and only if

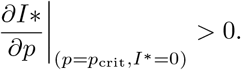

Notice that

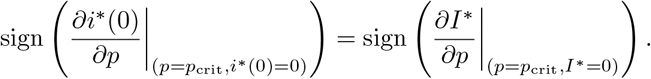

In our bifurcation analysis, we investigate the sign of the

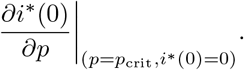

By differentiating the polynomial equation (21) implicitly with respect to parameter *p*, evaluating at *p* = *p*_crit_ and *i*^*\**^(0) = 0, and solving for *∂i*^*\**^(0)*/∂p* (at *p* = *p*_crit_ and *i*^*\**^(0) = 0), we obtain

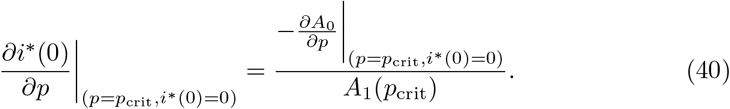

We now show that the right hand side of the above equality is positive for each parameter being investigated.

**Case 1 (***p* = *ρ*): First, we compute the partial derivative of coefficient *A*_0_ with respect to *ρ*:

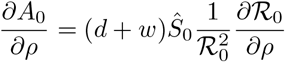

where

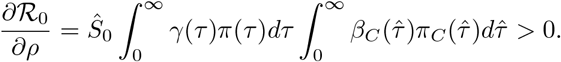

We then have

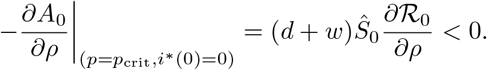

We now determine the sign of *A*_1_(*ρ*_crit_) (note through simplifications obtained on the numbers *χ* and 𝒴, coefficient *A*_1_’s dependence on parameter *ρ* is only through ℛ_0_). We have

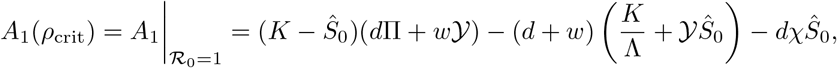

where Ŝ_0_ = *K*(1 − Λ*/d*). In replacing *Ŝ*_0_ with *K*(1 − Λ*/d*), we see that we can factor our *K*, so we analyze the sign of *A*_1_(*ρ*_crit_)*/K* to determine the sign of *A*_1_(*ρ*_crit_):

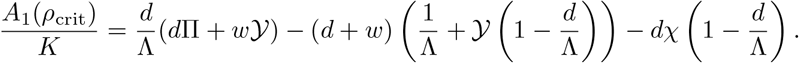

Additionally, we use Lemma 3.2 (specifically, we use 𝒴 = 1*/d*) to rewrite the above equation as follows:

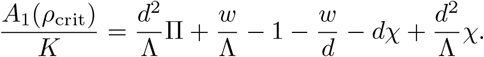

Since 𝒴 = Π + *χ* = 1*/d*, the first and last terms of the right-hand side of the above equation can be combined and re-expressed as *d/*Λ. We apply this simplification and rearrange terms to obtain

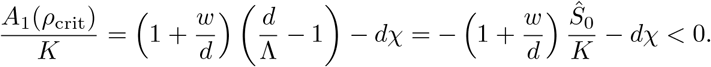

Thus, we have

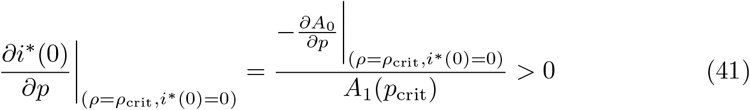

when *p* = *ρ*.

**Case 2 (***p* = *K*): First we compute the partial derivative of coefficient *A*_0_ with respect to *K* and evaluate at the critical parameter value:

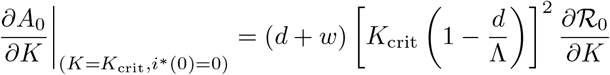

where

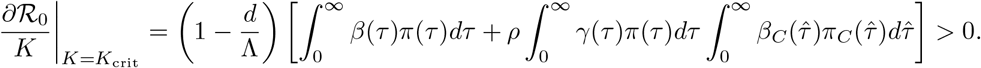

So

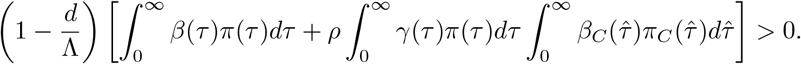

We determine the sign of *A*_1_(*K*_crit_) by observing

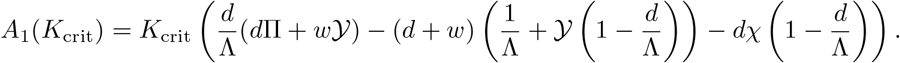

In Case 1, we already showed that

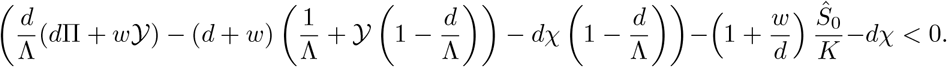

Thus, we have

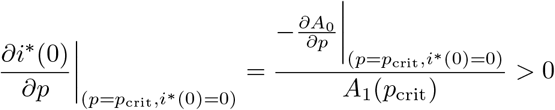

when *p* = *K*.

**Case 3 (***p* = Λ): First we compute the partial derivative of coefficient *A*_0_ with respect to Λ and evaluate at the critical parameter value:

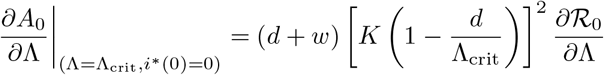

where

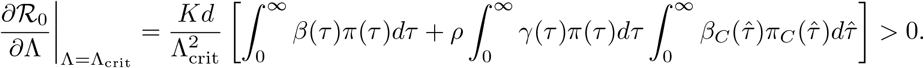

This implies that

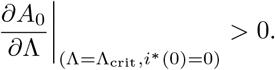

We determine that *A*_1_(Λ_crit_) is negative by observing

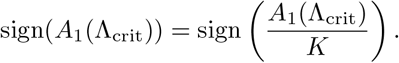

Through using simplifications that were used in Case 1, we obtain

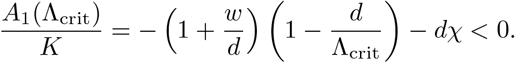

Thus, we have

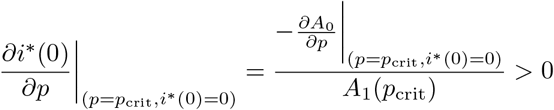

when *p* = Λ. This completes our proof.

Notice that bifurcation analysis with respect to the within-host parameters can only be done numerically since the epidemic parameters, including *β, β*_*C*_, *γ*, and *α* are formulated as functions of within-host variables where solutions for these immunological variables cannot be obtained explicitly. Furthermore, due to multiple logistic terms in system (8), analysis on the stability of the endemic equilibrium is not feasible (Gulbudak et al., 2017).

### 4 Simulations of the immuno-epidemiological model

We develop a finite d ifference sc heme co mbined wi th an OD E so lver in MATLAB in order to numerically solve the immuno-epidemiological model. We consider the within-host model of an infectious individual during the acute phase (1) with time-since-infection ranging from 0 ≤ *τ* ≤ *τ*_*end*_. The mesh chosen for the interval [0, *τ*_*end*_] can have equal step size ∆*τ* with *L* mesh-points. The output of the ODE solver is the solution vector, denoted here 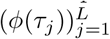 . We utilize MATLAB solver ode45, which adaptively chooses the time partition and interpolates at time points 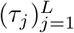 . Similarly, we consider the within-host model of a carrier infected individual (2) with time since becoming a carrier ranging from 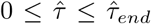. The mesh chosen for the interval 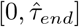 can have equal step size 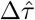 with 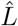 mesh-points. The output of the ODE solver is the solution vector, denoted here as 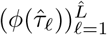 . For this model, we also utilize MATLAB’s solver ode45.

Moreover, in Table 3, the initial conditions for state variables *A*_*C*_, *I*_*C*_, and *P*_*C*_ are set to match the initial conditions for state variables *A, I*_*A*_, and *P* . That is, the solution of the within-carrier model would have a region for which the viral load in the bloodstream *P*_*C*_(*τ*) is positive. On the interval 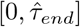, we find the time *τ*_∗_ for which *P*_*C*_(*τ*_∗_) *<* 10^−4^, and we assume that *τ*_∗_ is the start time of the carrier phase since this is the time at which the pathogen clears in the bloodstream. In our between-host simulations, we ignore time interval [0, *τ*_∗_). Note that we can set *τ*_*end*_ = *τ*_∗_ since this can represent the duration of the acute phase.

After obtaining solution vectors for systems (1) and (2), we can then discretize the linking functions. For this model we have linking functions corresponding to infected individuals during the acute stage, *f* (*τ*) ∈ *{β*(*τ*), *γ*(*τ*)*}*, and corresponding to infectious individuals during the carrier stage, 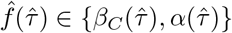. We can then generate a vector solution for each linking function corresponding to the acute infected, *f*_*j*_ = *f* (*τ*_*j*_), (with *j* = 1, …, *L*) and corresponding to the carrier infected 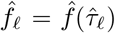 (with 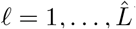). It is worth mentioning that for the simulations presented in Figure 6, linking functions 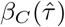 and 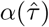 are set to 0 for all 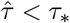 since the individuals are still in the acute phase.

For the epidemiological model, we approximate solutions to the age-structured PDE model with the stored within-host calculations by using a mixed implicit-explicit scheme. The numerical scheme is detailed in Appendix B.

#### 4.1 Parameter estimation

The parameter values used in the nested multi-scale model (1,2,8) are given in Table 3. The within-acute model (1) parameters comes directly from our prior work Macdonald et al. (2022). The least squares approach is used to fit the model to time-series data consisting of multiple needle-infected hosts during the acute stage of the disease (Perez-Martin et al., 2022). Monte Carlo simulations are also administered in (Macdonald et al., 2022) to assess identifiability and confidence in our fitting procedure. Some of these hosts may or may not have experienced a carrier stage of the disease (and such categorization is not taken into consideration during parameterization). In our simulations, we use the average fitted values that correspond to the buffalo who were needle-infected with the SAT1 strain. The fitted values here represent the average immune and viral kinetic dynamics of FMD infected buffalo during the acute phase. We fix the same values for the within-carrier model (2) but calibrate parameter values *K*_*T*_, *δ, η, v*, and *θ*_*C*_. We calibrate these parameters by using Mathematica’s manipulate feature to obtain a numerical solution of the ODE system (2) whose values align with the mean data values of four SAT1 needle-infected buffalo during the carrier stage (note these buffalo were hosts that were studied in the data that was used in Perez-Martin et al. (2022)). To prevent the innate immune response from being decoupled from the within-acute model, we fix *θ* as being the calibrated value that we obtained for *θ*_*C*_.

For calibrating the linking function parameters, parameter values are chosen to align with various quantities presented in Jolles et al. (2021). They estimate the infectious period for an acute phase as being 5.7 days and the duration of the carrier phase as being 243 days. It is worth noting that the duration of the carrier stage can be longer; it can even last for years (Stenfeldt C, 2020). In Jolles et al. (2021), they approximate the basic reproduction number in the absence of carriers as being 15.8 and the basic reproduction number with carriers as being 23.8. Additionally, they find that the transmission rate due to contacting an infectious individual during the acute phase as being two orders of magnitude larger than the order of magnitude of the transmission rate corresponding to coming in contact with a carrier-infected individual.

After storing solution vectors for system (1), we can approximate ℛ_*A*0_ by using the trapezoidal rule to approximate the first integral given in (11) over the interval [0, *τ*_*end*_]. Now, the stored vectors used for the carrier system (2) uses initial values set to align with the beginning of the acute phase, so for approximating ℛ_*C*0_, we use the trapezoid rule to approximate the second integral given in (11) over the interval 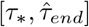 where *τ*_∗_ is the time in which *P*_*C*_(*τ*_∗_) *<* 10^−4^ (which estimates the start of the carrier phase). Using these methods and the calibrated parameter values described in Table 3, we have that ℛ_*A*0_ ≈ 15.6426, ℛ_*C*0_ ≈ 7.3761, and ℛ_0_ ≈ 23.9187.

In subfigure 5a, we have that an infected individual experiences an increased probability of pathogen clearance in the bloodstream during days 5-8, and is over 99% likely to have pathogen clearance in the bloodstream by 7.6 days after infection. In subfigure 5b, we present a graph of the function that represents the probability of pathogen clearance in the tonsils 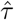 days after becoming a carrier. From this plot, we have that a carrier-infected individual is about 76.68% likely to recover by 243 days (after becoming a carrier) and over 99% likely to recover by day 395 days (after becoming a carrier). In Figure 5, we have plots of the transmission rates *β*(*τ*) and 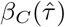. Observe that the peak value of the function *β*(*τ*) is two orders of magnitude larger than the peak value of 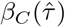, which aligns with what was observed in Jolles et al. (2021). Epidemiological parameters *w, ρ*, and *d* are chosen to align with what was used in Jolles et al. (2021) while the remaining parameters, the birth rate Λ and the carrying capacity *K*, are adjusted to match ℛ_0_.

**Fig. 5:**
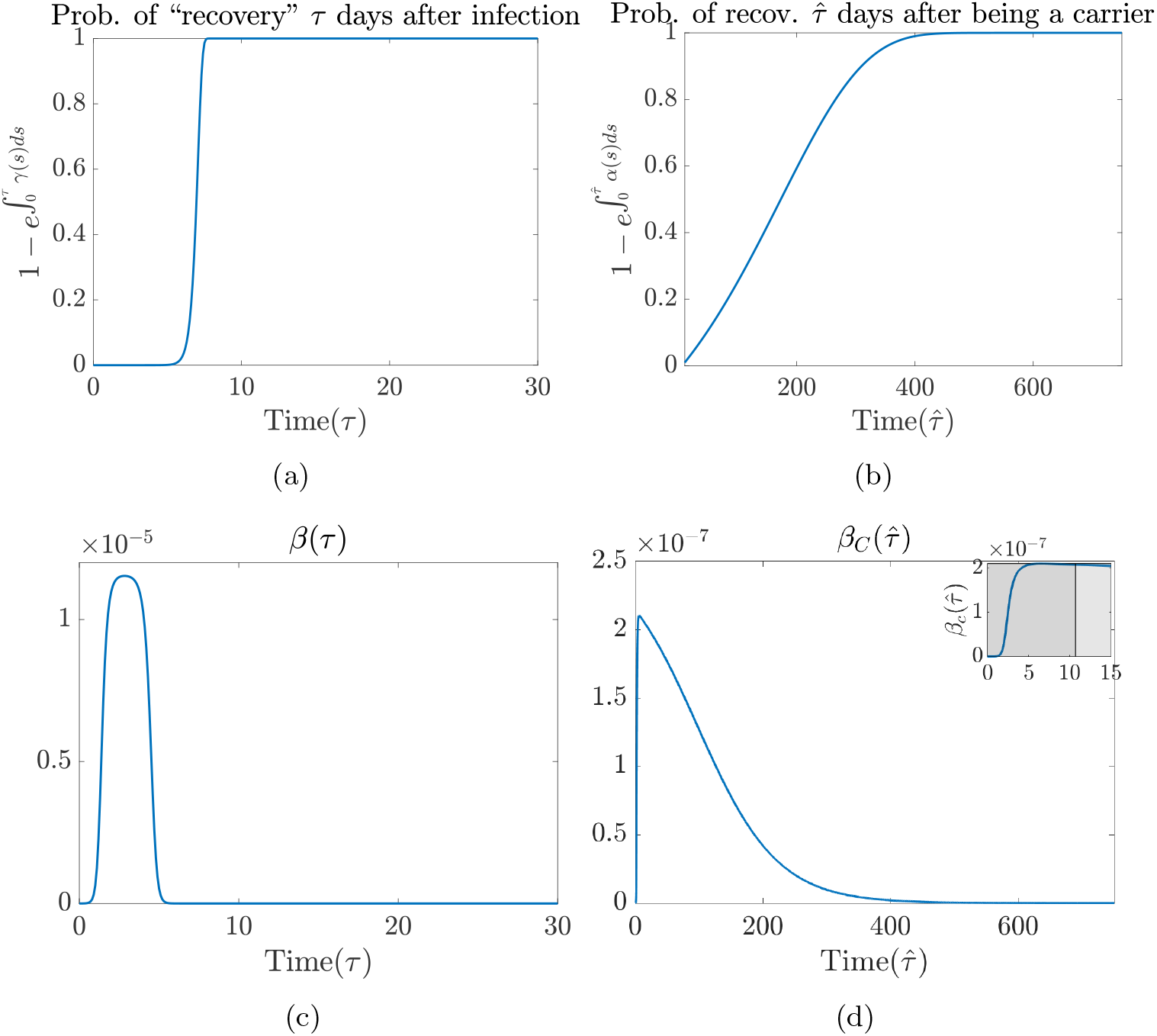
Top left. Subfigure displays the probability of pathogen clearance in the bloodstream. **Top right**. Subfigure displays the probability of pathogen clearance in the tonsils by day 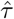 (after becoming a carrier). **Bottom left**. Subfigure displays the transmission rate *β*(*τ*) due to interacting with an acute infected individual. **Bottom right**. Subfigure displays the transmission rate 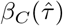 due to interacting with a carrier over time interval [0, 750]. The shaded inset figure shown in subfigure (d) illustrates that the start of the carrier phase *τ*_*\**_ = 10.7. In our simulations for the nested multi-scale model, we ignore the within-host carrier dynamics on interval [0, *τ*_*\**_), by setting 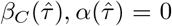 for all 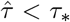. The linking function parameter values used are given in Table 3.

**Fig. 6:**
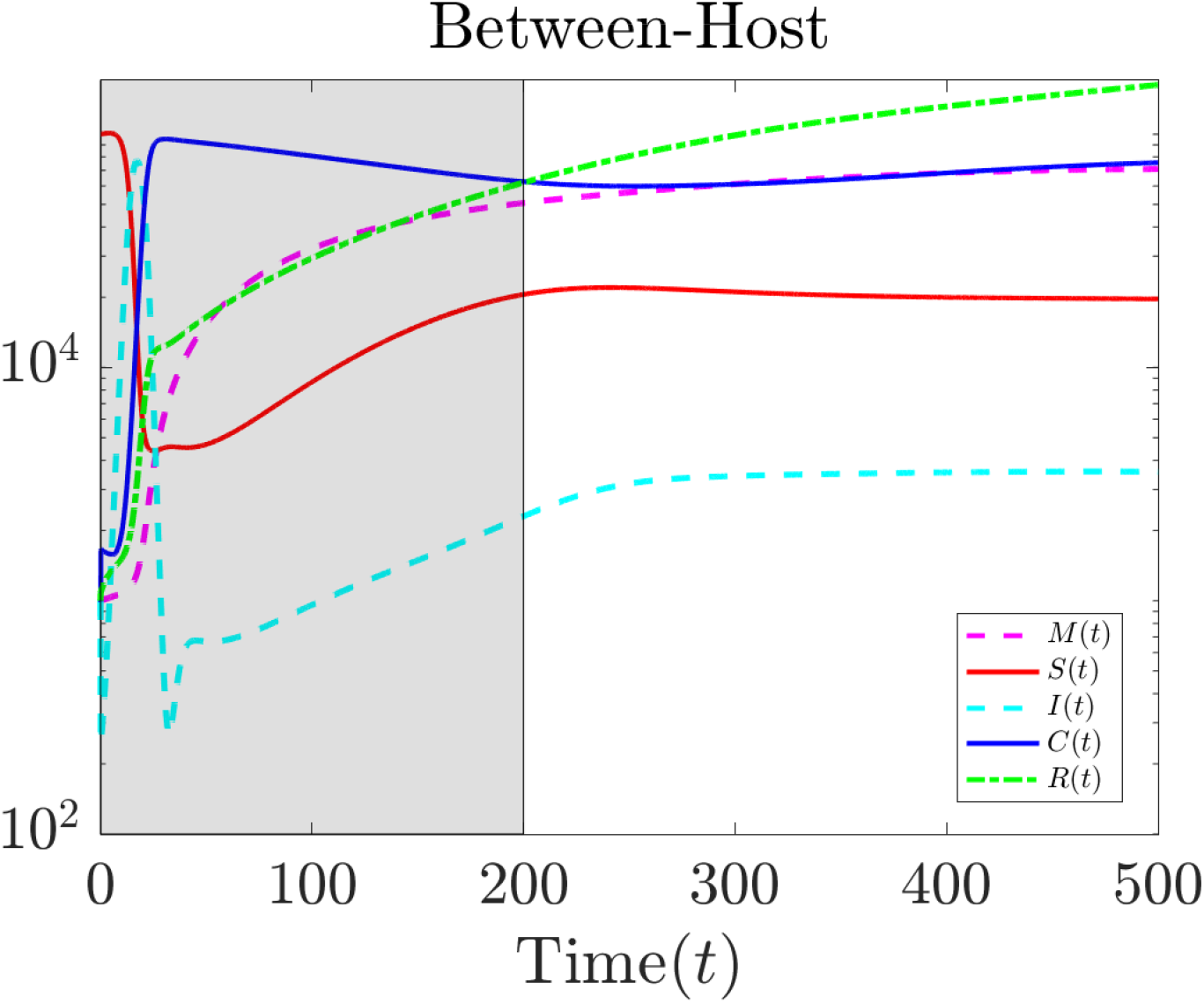
Logarithmic plots of model compartments *M* (*t*), *S*(*t*), *I*(*t*), *C*(*t*), and *R*(*t*). The shaded region shows that the susceptible population does not fade out due to new births. New susceptible individuals and the presence of carrier-infected individuals lead to a re-emergence of acute-infected individuals.

#### 4.2 Numerical results and epidemiological implications

In Figure 6, we have logarithmic simulations of the classes *M* (*t*), *S*(*t*), *I*(*t*), *C*(*t*), and *R*(*t*) during the first 500 days, and in Appendix D, we present Figure 12 (and Figure 13), which gives the logarithmic plots (and standard plots) of each compartment and the corresponding endemic steady state solution over the span of 2000 days to illustrate that the numerical solution to our model is approaching the endemic equilibrium.

As shown in Figure 6, we see that the replenishing of new susceptible individuals (red) and the presence of carrier-infected individuals (blue) are the driving factors for FMD disease persistence. These results (which are illustrated in the gray region) align with what was observed in Jolles et al. (2021). They considered two mechanisms for persistence of FMDV: (1) FMDV might be maintained as “typical childhood infections” do, where the elder born calves of one year spark an epidemic in the earliest born of the following cohorts, and (2) persistently-infected buffalo are present in the population and continue to infect other individuals. In the gray shaded region of Figure 6, we have the maternally immune (magenta and dashed) steadily increasing due to new births. The susceptible population (which began with 100,000 individuals) drops to being as low as 4,282 individuals by day 26. This leads to the acute infected population (in the cyan and dashed plot line) peaking at over 72,900 individuals by day 16. By day 30, the carrier-infectious class peaks at more than 95,000 individuals. By day 33, the acute-infected class drops to as low as 270 individuals, and this is largely due to how many acute-infected individuals quickly transition to either class *C*(*t*) or *R*(*t*) (green). Class *C*(*t*) does not drop as significantly as class *I*(*t*). This is because the carrier stage is significantly longer than the acute stage. From days 26 to 50, we do not see the susceptible population fading out but rather remaining somewhere near 4,282 to 4,682 individuals. This phenomenon is likely due to births of new susceptible individuals and/or due to individuals in class *M* (*t*) entering class *S*(*t*) through waning immunity. From days 50 to 200, we observe a steady increase in the numbers of susceptible individuals, and also a similar rate of increase in class *I*(*t*) at this time. By day 250, there are 3,170 acute infected individuals, and we do not see any significant declines in class *I*(*t*) during time interval [250, 500].

### 5 Impact of within-host parameters on ℛ_0_

Sensitivity analysis (SA) is a tool that helps one quantifiably assess the impact of changes in model parameters on infection dynamics, which can help one measure the expected efficacy of disease control strategies. In this section, we use an SA method for multi-scale models developed in our prior work Gulbudak et al. (2022) to investigate the impact of within-host parameters on ℛ_0_.

The elasticity or normalized forward sensitivity index of a quantity of interest (QOI) *q* to a parameter of interest (POI) *p* can be defined as the ratio of the relative change in the variable to the relative change in the parameter:

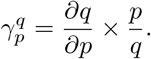

In our case, *q* is the basic reproduction number ℛ_0_ for the original system (8), *p* is any within-host viral and immune kinetics parameter during either the acute or carrier phase.

In our numerical experiments, we compute the sensitivity index 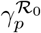 when a within-host parameter *p* is set to a value around a baseline parameter value (see details in Appendix C). The parameters of interest and the ranges of parameters are given in Table 5. We rank the sensitivity index at the baseline values in Table 6 and observe the following:

**Table 5:** Ranges for parameters used in the sensitivity analysis. Note parameter *a* is only being varied in the within-carrier model.

| Parameter | Sensitivity Analysis Range |
| --- | --- |
| Parameters for acute infectious |  |
| $b$ | [0.09, 1] |
| $K_A$ | [1, 12] |
| $r_A$ | [1, 10] |
| $\theta$ | [$10^{-6}$, 0.1] |
| $\mu$ | [$10^{-6}$, 0.2] |
| $\xi$ | [0.1, 4] |
| $\psi$ | [$10^{-2}$, 0.7] |
| $\Omega$ | [0.1/21.23, 2/21.23] |
| $\omega$ | [$10^{-3}$, 0.5] |
| Parameters for carriers |  |
| $a$ | [0, 0.1] |
| $b_C$ | [0.09, 1] |
| $K_C$ | [1, 12] |
| $K_T$ | [1, 12] |
| $r_C$ | [1, 10] |
| $\delta$ | [0.00005, 0.017] |
| $\eta$ | [$10^{-5}$, 0.1] |
| $\theta_C$ | [$10^{-6}$, 0.1] |
| $\mu_C$ | [$10^{-6}$, 0.2] |
| $\nu$ | [0.0025, 5.25] |
| $\xi_C$ | [0.1, 4] |
| $\psi_C$ | [$10^{-2}$, 0.7] |
| $\Omega_C$ | [0.1/21.23, 2/21.23] |
| $\omega_C$ | [$10^{-3}$, 0.5] |

**Table 6:** Parameters and their sensitivity index ranked from most to least impactful. Note that the sensitivity index 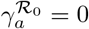 because our baseline value for *a* is 0.

| Rank | Parameter | $\gamma_p^{\mathcal{R}_0}$ | Rank | Parameter | $\gamma_p^{\mathcal{R}_0}$ |
| --- | --- | --- | --- | --- | --- |
| 1 | $b$ | -0.653 | 13 | $\theta$ | 0.00117 |
| 2 | $K_T$ | 0.338 | 14 | $\theta_C$ | $7.43 \cdot 10^{-4}$ |
| 3 | $\eta$ | -0.307 | 15 | $\psi_C$ | $5.88 \cdot 10^{-4}$ |
| 4 | $K_A$ | -0.294 | 16 | $\psi$ | $3.39 \cdot 10^{-4}$ |
| 5 | $r_C$ | -0.0843 | 17 | $\mu_C$ | $-1.93 \cdot 10^{-4}$ |
| 6 | $b_C$ | -0.0781 | 18 | $\mu$ | $-1.15 \cdot 10^{-4}$ |
| 7 | $\nu$ | 0.0747 | 19 | $\Omega_C$ | $4.87 \cdot 10^{-5}$ |
| 8 | $\xi_C$ | 0.0712 | 20 | $\Omega$ | $3.61 \cdot 10^{-5}$ |
| 9 | $\xi$ | -0.0704 | 21 | $\omega$ | $-2.04 \cdot 10^{-5}$ |
| 10 | $K_C$ | -0.0408 | 22 | $\omega_C$ | $-1.94 \cdot 10^{-5}$ |
| 11 | $r_A$ | 0.00848 | 23 | $a$ | 0 |
| 12 | $\delta$ | 0.0071 | | | |

- All parameters corresponding to the innate immune response (at baseline values) had the least impact on ℛ_0_. These results suggest that the innate immune response’s performance on targeting the pathogen is least efficacious.
- At the baseline (estimated) values of the within-host immune parameters for both the acute and carrier stage, the within-host adaptive response activation rate during the acute phase *b* (with baseline value being 0.37) has the largest impact on ℛ_0_: a 1% increase in *b* causes a 0.653% reduction in *R*_0_. This suggests that the most efficient control measures are the ones that boost the adaptive immune response.
- The within-host pathogen carrying capacity in the tonsils during the carrier phase, *K*_*T*_ (with baseline parameter value 8) has the second largest impact on ℛ_0_: a 1% increase in *K*_*T*_ causes a 0.338% increase in ℛ_0_.
- The decay rate of the pathogen in the tonsils, *η*, at the baseline parameter value 0.008 has the third largest impact on ℛ_0_: a 1% increase in *η* causes a 0.307% reduction in ℛ_0_.

In Figures 7 and 8, we provide plots of the sensitivity index (in red) when a within-host parameter *p* is varied and plots of the resulting *R*_0_ (in blue and with baseline value starred). Note that Figure 7 is organized to have the subfigures on the left-hand side correspond to the within-acute parameters, while the right subfigures correspond to the within-carrier parameters. The figures corresponding to the impact of the innate immune response parameters (except for parameter *a*) on *R*_0_ are given in Appendix D (see Figure 14) since they were the least impacting on *R*_0_. From an immunological standpoint, it makes sense for the innate response parameters to not be as effective in changing *R*_0_ in comparison to the adaptive immune response since the primary role of the innate immune response is to send alarm signals that lead to the production of the adaptive immune response.

**Fig. 7:**
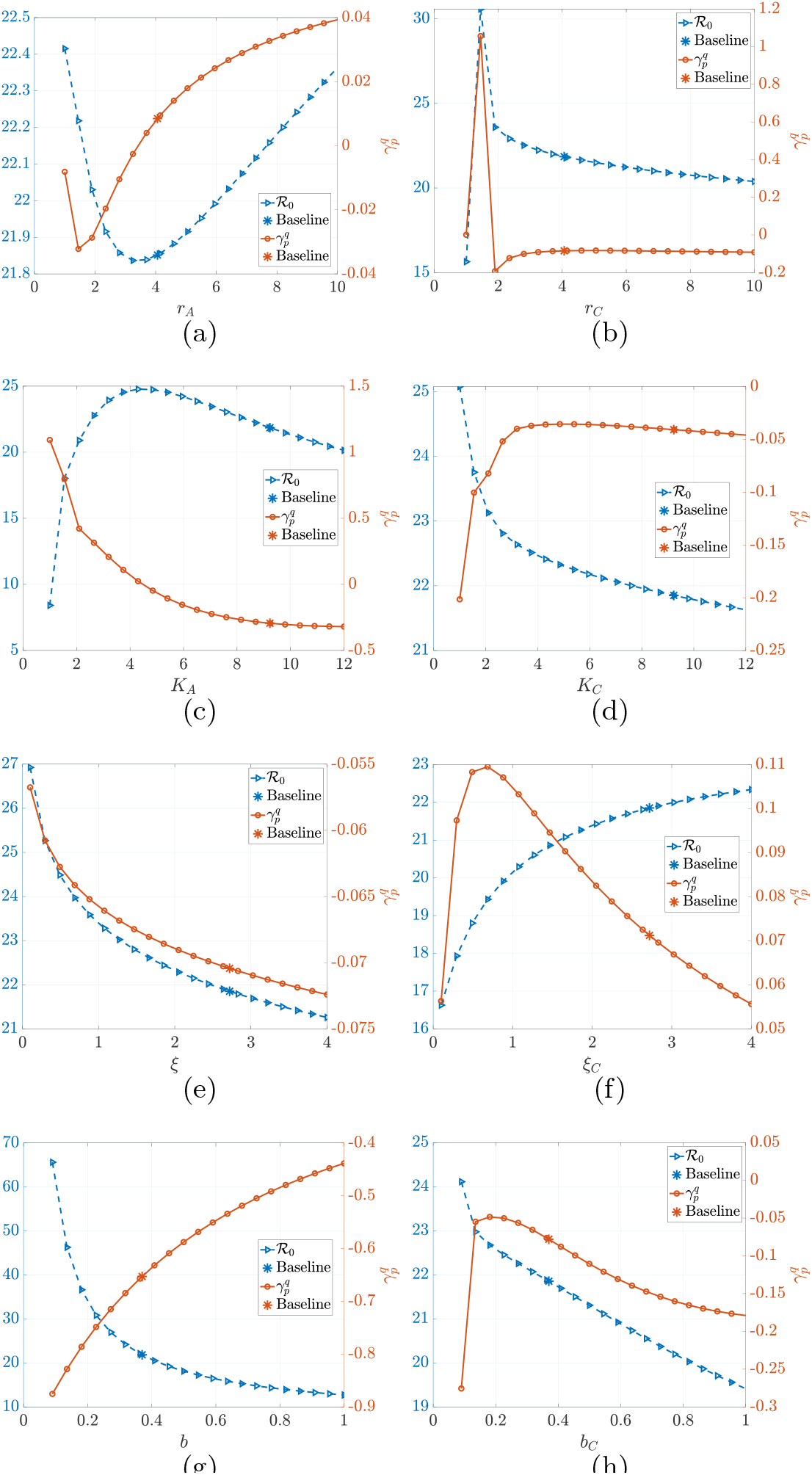
Impact of viral and immunological parameters (*r*_*A*_, *r*_*C*_, *K*_*A*_, *K*_*C*_,, *ξ, ξ*_*C*_, *b*, and *b*_*C*_) on *R*_0_. The virus-immune kinetics-dependent epidemiological reproduction number, ℛ_0_ (left y-axis, blue curve), along with the response curves for the extended SA, 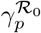 (right y-axis, red curve), are plotted against the immune parameter values *p* over a range of values (see Table 5) around the baseline estimates (see Table 3).

**Fig. 8:**
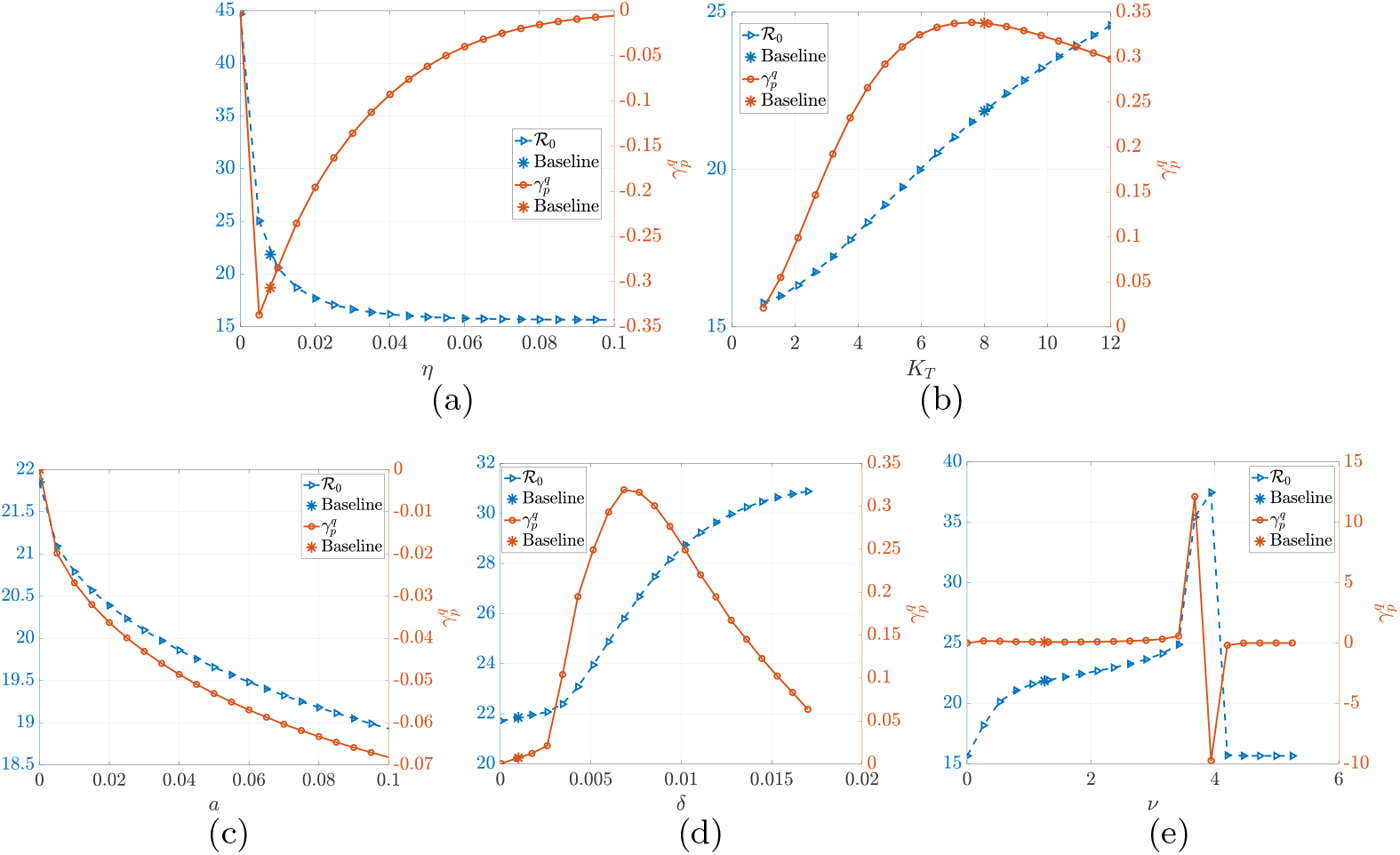
Impact of within-carrier parameters *ν, η, K*_*T*_, *a, δ* on ℛ_0_. The immune-response-dependent epidemiological reproduction number, ℛ_0_ (left y-axis, blue curve), along with the response curves for the extended SA, 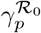(right y-axis, red curve), are plotted against the immune parameter values *p* over a range of values (see Table 5) around the baseline estimates (see Table 3). In subfigure (e) the SA results for *ν* exhibit unusual behavior when *v* ∈ [3.41337, 4.2005]. In subfigure 9a, we rerun sensitivity analysis results for *ν* but over a finer partition of [3.41337, 4.2005].

In subfigure 7 g, we also observe that increasing the adaptive immune response activation rate, *b*, during the acute stage leads to a reduction in ℛ_0_. As mentioned above, the parameter *b* has the largest impact on ℛ_0_ at the baseline parameter values.

Our SA results corresponding to parameter *b* align with our prior analysis on the acute stage of the disease (Macdonald et al., 2022). In our prior work, we observed a strong correlation between the adaptive immune response activation rate and the acute infection time period. We find this correlation to be one of the key influences on ℛ_0_. In subfigure 7h, we observe that increasing the adaptive immune response activation rate during the carrier phase, *b*_*C*_, also leads to a decrease in ℛ_0_. Moreover, the parameter *b*_*C*_ is ranked as having the sixth largest impact on ℛ_0_ at the estimated baseline values. Since the adaptive response activation rate during the carrier phase has an impact on the reduction of ℛ_0_, continual application of a control that boosts the adaptive immune response during the carrier stage should have an epidemiological impact on reducing the spread.

The rankings of impact of the within-host carrying capacity in tonsils *K*_*T*_ and the decay rate of the pathogen in the tonsils *η* suggest that the pathogen load in the tonsils plays a crucial role at the epidemiological scale and control measures that kill the virus in the tonsils should be investigated. It is worth noting that such a result may be a direct consequence of the formulation of the linking functions 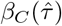 and 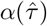, which should be validated with data.

In subfigures 7a-7b, we have the sensitivity index of the viral replication rate in the blood during the acute phase *r*_*A*_ and the sensitivity index of the viral replication rate in the blood during the carrier phase *r*_*C*_ . In our SA results, we numerically observe that an increase in *r*_*A*_ leads to an increase in ℛ_0_. In our prior work Macdonald et al. (2022), we observed that the viral growth rate has a negative correlation with the latent period in that buffalo having fast-growing viral populations transfer to the infectious class more rapidly. As mentioned in other works (Grassly and Fraser, 2008; Sartwell, 1995; Fine, 2003) short incubation periods play a significant role in the rapid spread of highly transmissible pathogens. This could explain why the viral growth rate has a significant impact on ℛ_0_.

An interesting result observed is that increases in *r*_*C*_ lead to decreases in ℛ_0_. This suggests that a higher viral replication rate could contribute to a shorter carrier stage. This can occur because increasing the viral growth rate can result in a shorter duration for which the pathogen remains in the bloodstream (due to increased levels in the adaptive immune response). The shorter the duration of the viral load in the blood-stream, the fewer opportunities for the pathogen to migrate to the tonsils. However, the SA results corresponding to varying *r*_*C*_ are only measuring how such a parameter change influences ℛ_0_ v ia i nfluencing ℛ_*C* 0_. Th e re sults pr esented in su bfigure 7a indicate a significant d rawback o f w hat h appens w hen a h igh v iral r eplication rate occurs during the acute phase. More specifically the higher the viral growth rate, the higher the transmission level during the acute phase. It is worth mentioning that in the within-carrier model (2) the pathogen migration term *v* can be viewed as a viral replication rate in the tonsils. In Table 6, we see that at the baseline parameter values, *v* behaves similarly to the viral replication rate in the bloodstream during the acute phase, *r*_*A*_, in that an increase in *ν* leads to an increase in ℛ_0_, but *v* has a higher impact in comparison to *r*_*A*_.

In subfigure 8e, the graphs of the computed sensitivity index and ℛ _0_ appear to be nonsmooth when *v* ∈ [3.41337, 4.2005]. In particular, when *ν* = 3.67575 the sensitivity index 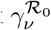 jumps to 12.12 and ℛ_0_ increases to 35.4174. When *v* = 3.93812, 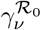drops to −9.74427 but ℛ_0_ = 37.467. The sensitivity index being negative here implies that *R*_0_ is decreasing near *v* = 3.93812. We can then infer that ℛ_0_ peaks at some value *v* ∈ (3.67575, 3.93812). We support this inference by rerunning the sensitivity analysis simulations for *ν* over a finer partition of [3.41337, 4.2005], and the results are given in subfigure 9a. The graph of the computed ℛ _0_ given in subfigure 9a is smooth, but the graph of the sensitivity index does not look smooth near *ν* = 4.04307. Note that using a finer p artition d oes n ot i mprove t his c haracteristic. T his i s b ecause t here i s some *v* ∈ (4.04307, 4.08243), for which the pathogen-free equilibrium *ε*_0_ (expression given in Table 2) is changing from being unstable to being locally asymptotically stable (Recall that at the baseline parameter values *ε*_0_ is unstable.). The immunological reproduction number ℛ_0_ (3) is approximately 1.00307 and 0.99341 when *v* = 4.04307 and *v* = 4.08243 (respectively). In Figure 9 we numerically solved for the within-carrier model (2) for varying values of *v* ∈ *{*3.96436, 4.00372, 4.04307, 4.08243*}*. When *v* = 3.96436, we observe viral relapse in the tonsils within the first 750 days. Because the transmission rate 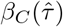 depends on the pathogen load in the tonsils, these instances of viral relapse lead to a larger value for ℛ_0_. As we increase *ν* to 4.00372 and 4.04307, we see the oscillatory behavior in the pathogen load for the tonsils dissipate and a decrease in ℛ_0_ is observed. Lastly, when *v* = 4.08243, we see that the solution is converging to *ε*_0_, and ℛ_0_ = 16.2211.

**Fig. 9:**
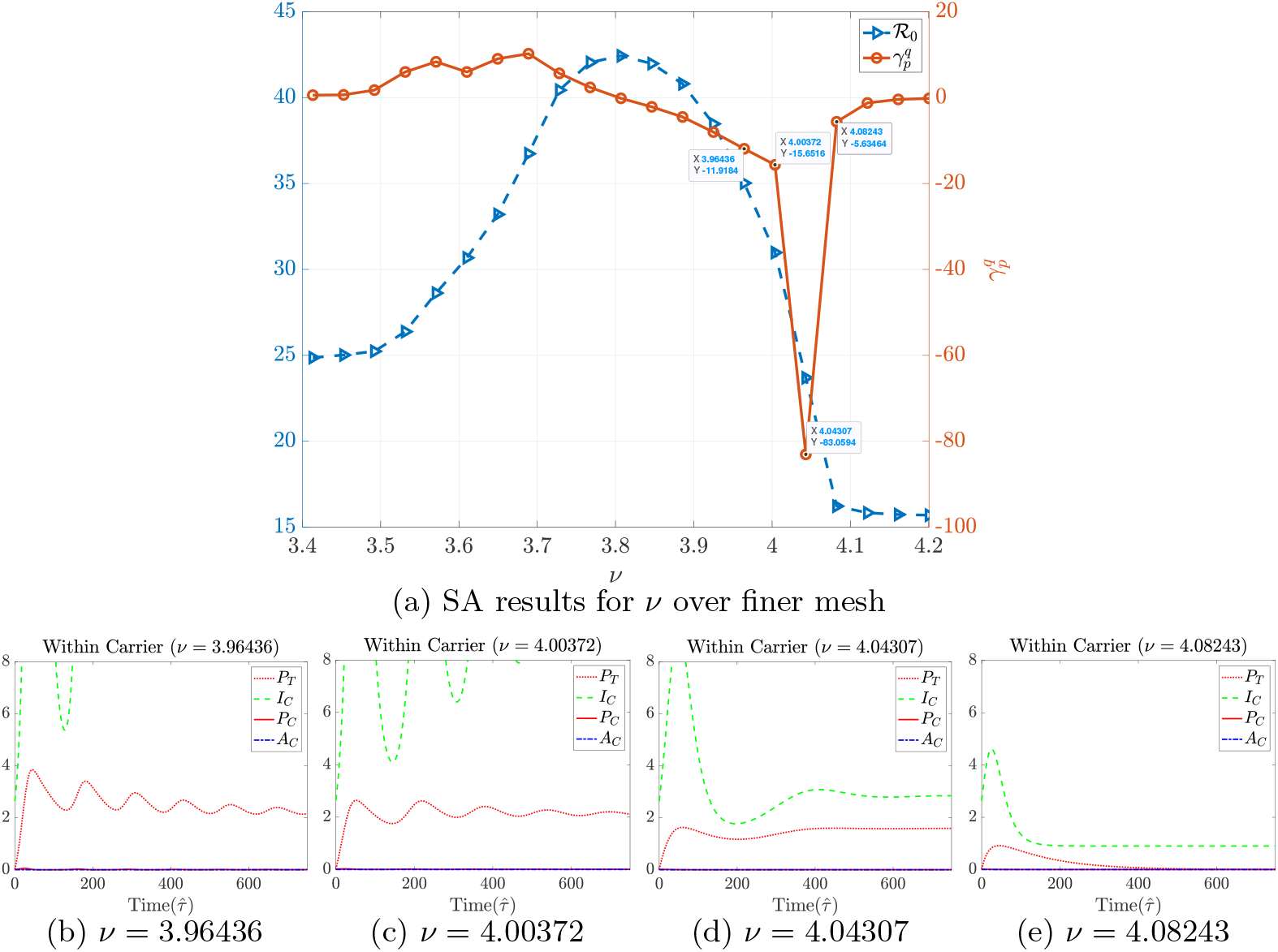
Sensitivity analysis results of *v* alongside the within-carrier solutions. In sub-figure ( a), w e p resent t he sensitivity analysis results t o s how that *R* _0_ i s smooth on interval *v ∈* [3.41337, 4.2005]. There is a dip in the sensitivity index at *v* = 4.04307. In subfigures ( b)-(e), we display solutions t o t he within-carrier model ( 2) with *v ∈ {*3.96436, 4.00372, 4.04307, 4.08243*}* and all other parameters set as described in Table 3. When *v* = 3.96436, 4.00372, and 4.04307 the solutions are converging to the the pathogen-persisting interior equilibrium *E*_2_, and when *v* = 4.08243 the solution is converging to the pathogen-free equilibrium *E*_0_ (See Appendix D Figure 15 which displays the same within-carrier simulations but over a longer time period to illustrate the convergence). The dip in the sensitivity analysis results corresponds to the immunological basic reproduction number switching from being ℛ_0_ *>* 1 to ℛ_0_ *<* 1. Observe also the computed epidemiological ℛ_0_ when *v* = 4.08243 is close to ℛ_*A*0_.

Our results presented in Figure 10 indicate that the carrier model (2) is more likely to display viral relapse in comparison to the acute model (1). Changes in the within-carrier parameter values can have an impact on immunological quantities (i.e., ℛ_0_, ℛ_1_, and ℛ_2_, which are given in Table 1) that determine the stability of a steady state solution to system (2). The duration of the carrier phase for an FMD infected-buffalo varies from months to years (Stenfeldt C, 2020; Shikumwifa, 2022). Instances of viral relapse could explain why some buffalo undergo a longer duration of the carrier phase in the sense that viral relapse would delay the process in clearing the pathogen in the tonsils. In the context of immunology, instances of viral relapse could arise in a carrier-infected buffalo i f s ome o f t he v iral p articles w ithin t he t onsils w ere t o drip back into the bloodstream; however, to the best of our knowledge no field studies are investigating such a phenomenon.

**Fig. 10:**
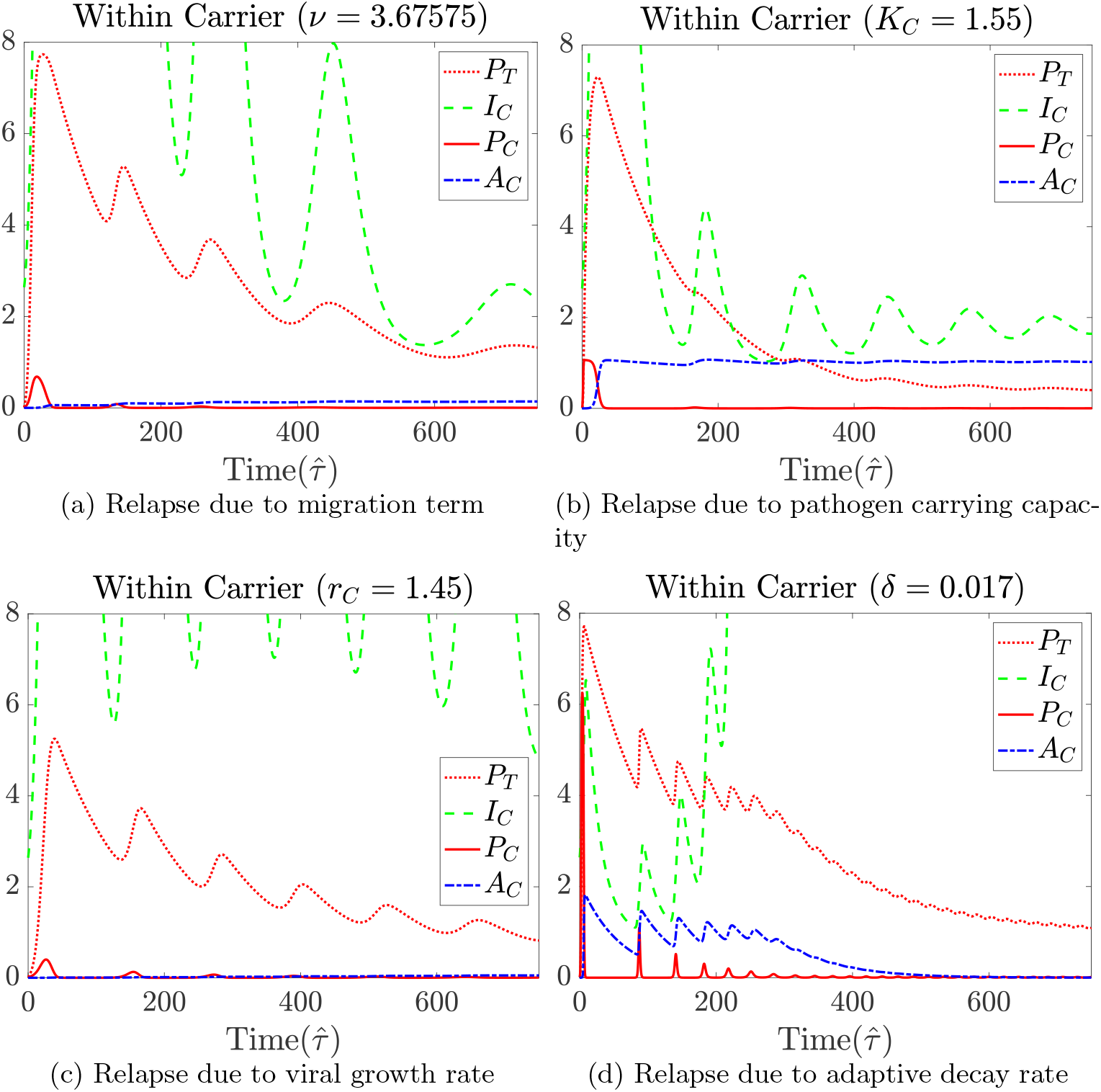
Instances of viral relapse from the carrier model (2). These subfigures demonstrate how the within-carrier model can display viral relapse. For each simulation, one of the within-carrier parameters was set to a different value, as described in the sub-figure titles, while the remaining within-carrier parameters were set to the baseline parameter values (see Table 3).

**Fig. 11:**
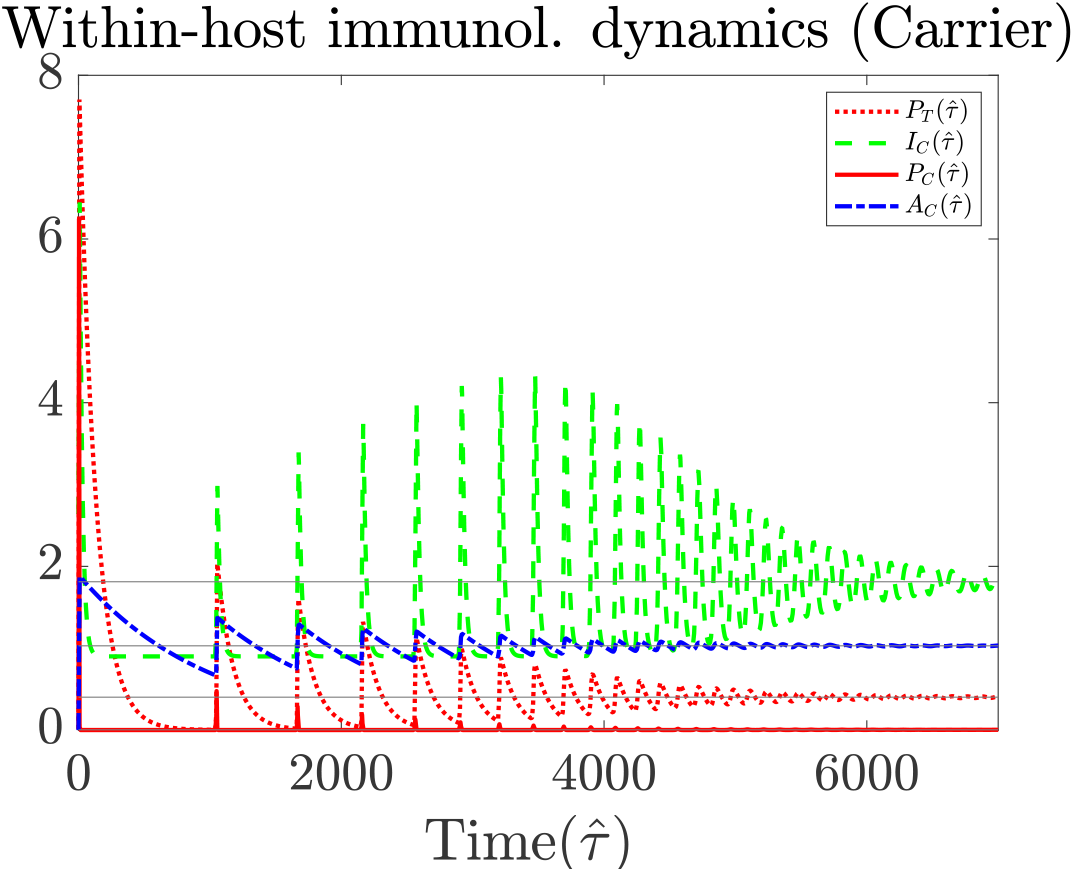
The within-carrier host dynamics over 7000 days to further illustrate how the solution to the within-carrier model is converging to the pathogen-persisting interior equilibrium.

**Fig. 12:**
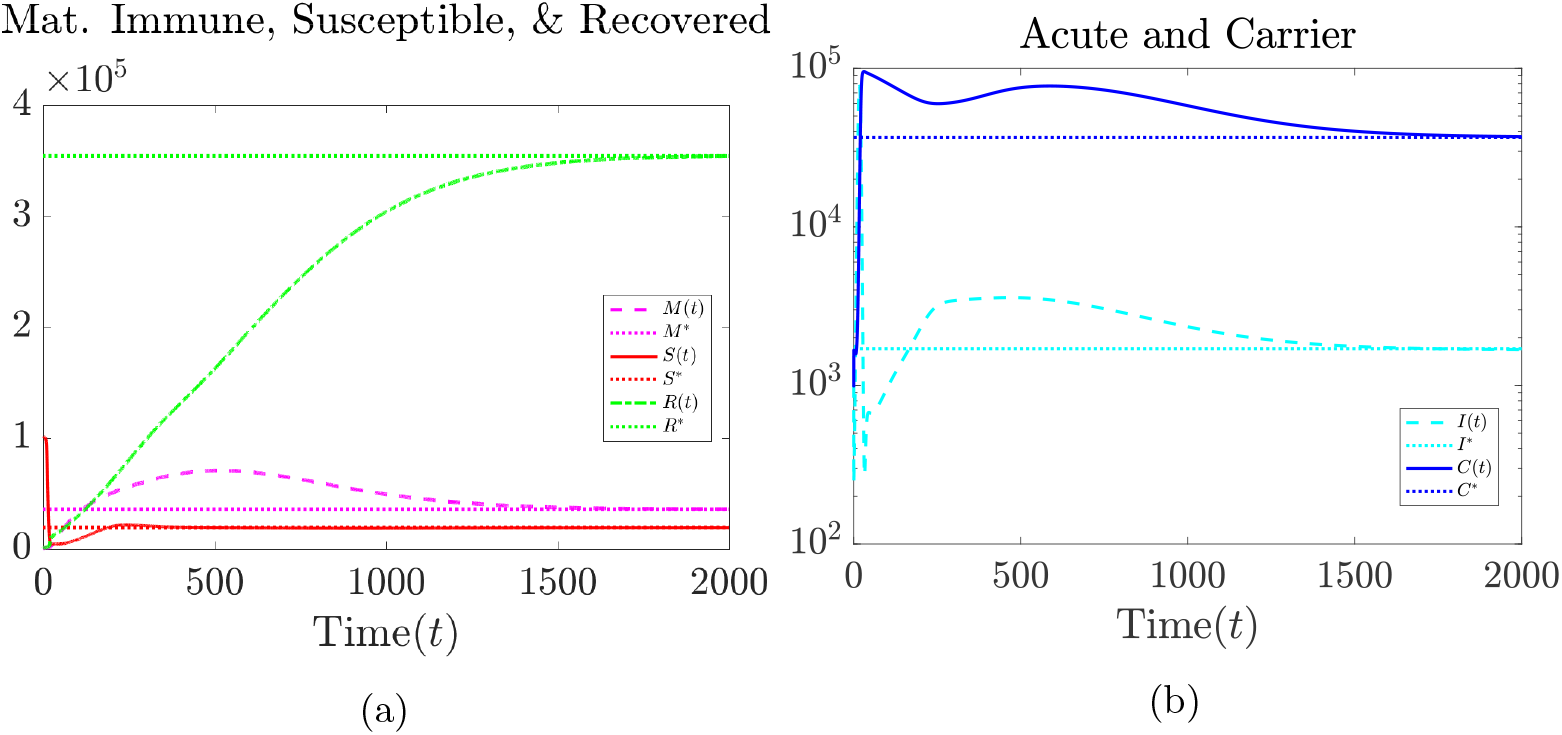
Between-Host Plots. **Left**. Logarithmic plots of classes *M* (*t*), *S*(*t*), and *R*(*t*), along with the steady states. **Right**. Logarithmic plots of classes *I*(*t*) and *C*(*t*), along with a plot of the numbers 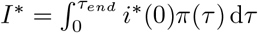and 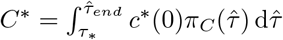. All parameters used are given in Table 3.

**Fig. 13:**
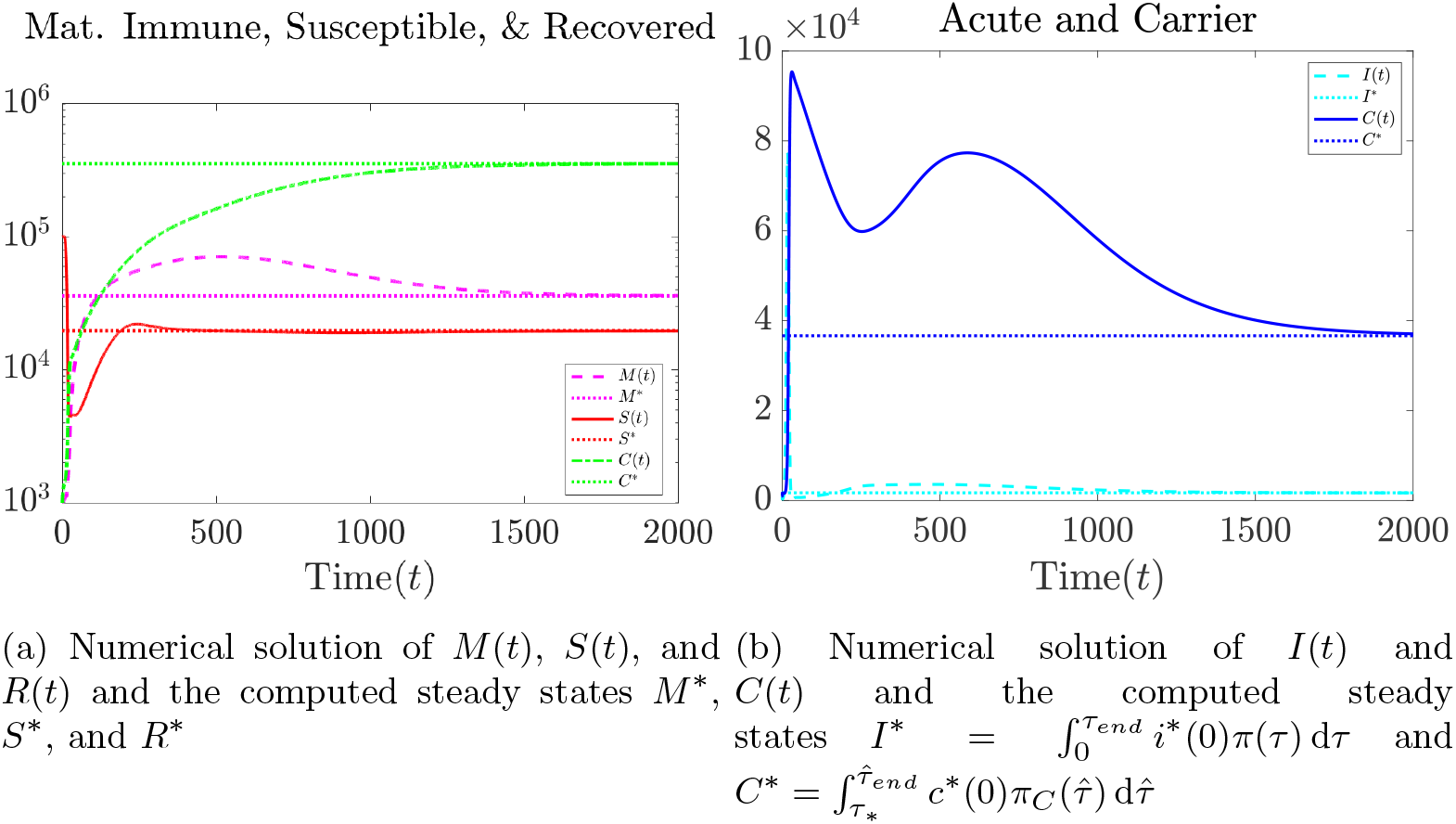
Between-Host Plots along with steady states. **Left**. Plots of classes *M* (*t*), *S*(*t*), and *R*(*t*). **Right**. Plots of classes *I*(*t*) and *C*(*t*). All parameter values used are given in Table 3.

**Fig. 14:**
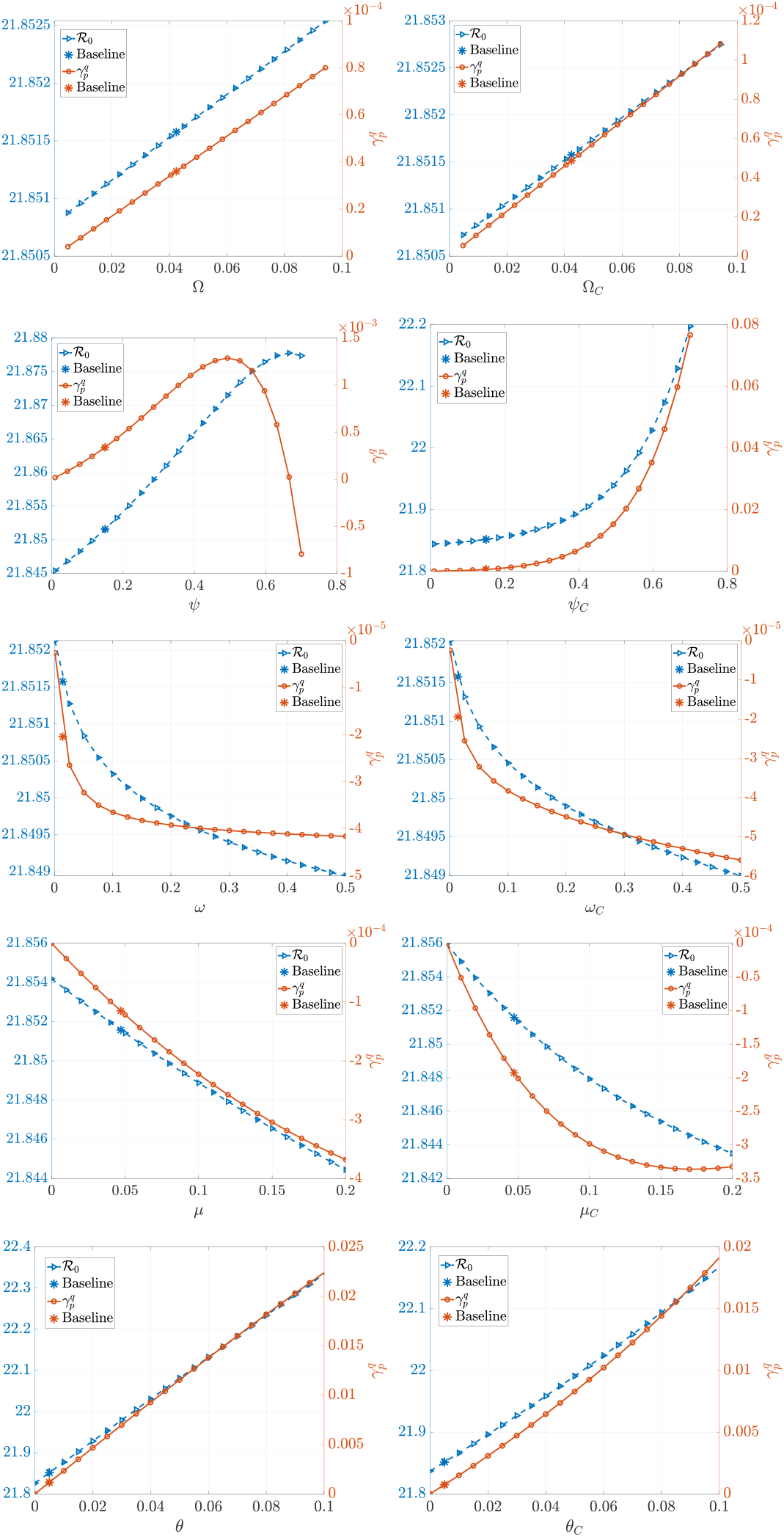
Sensitivity analysis results. The figures show the impact of the innate immune response parameters (Ω, Ω_*C*_, *ψ, ψ*_*C*_, *ω, ω*_*C*_,*μ, μ*_*C*_, *θ, θ*_*C*_) during the acute and carrier stage.

**Fig. 15:**
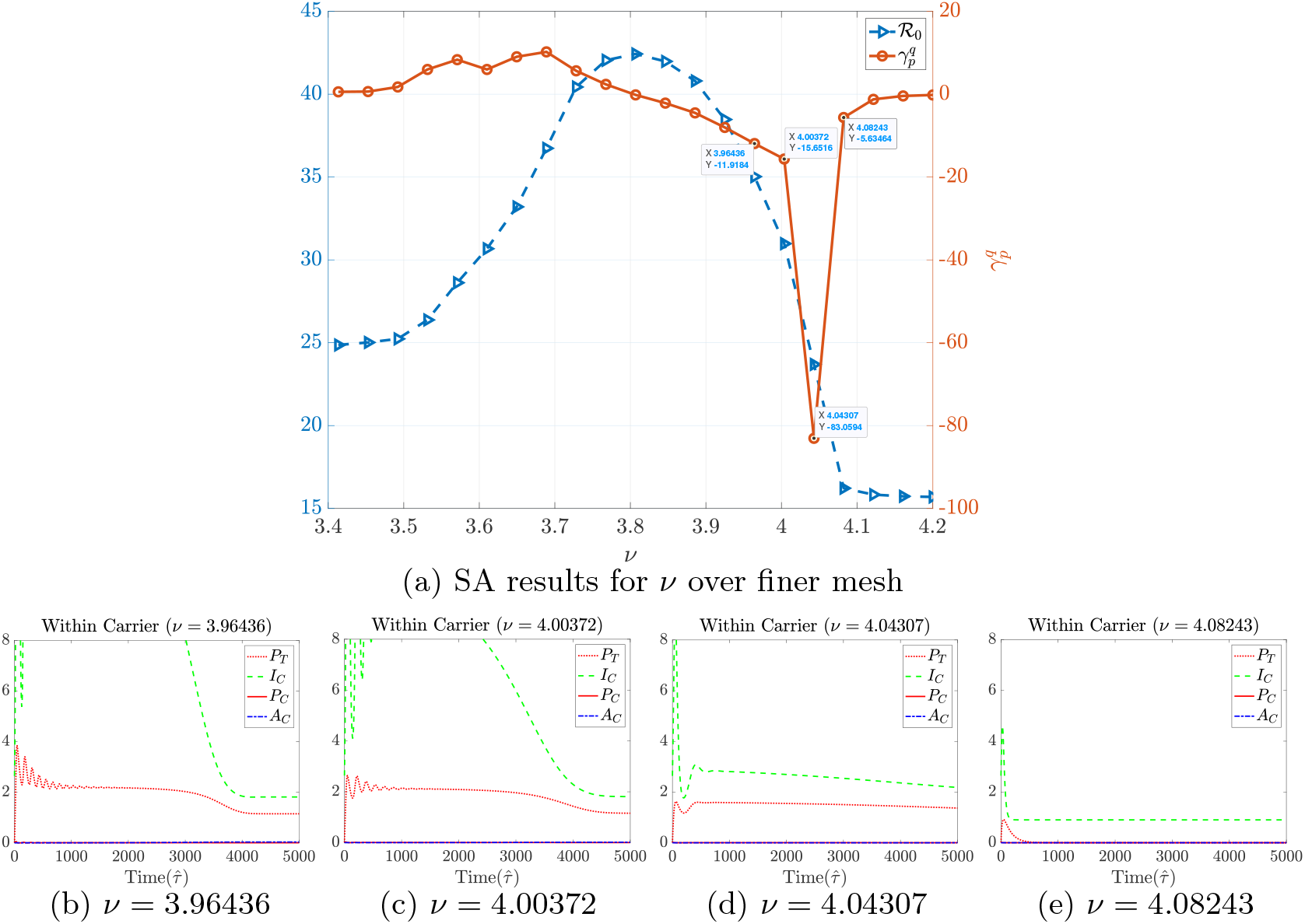
Sensitivity analysis results of *v* alongside the within-carrier solutions to model (2) as described in Figure 9 but displayed over a longer time period. The within-carrier solutions in Figures (b)-(d) are converging to the pathogen-persisting equilibrium the solution in Figure (e) is converging to the pathogen-free equilibrium.

### 6 Discussion

Foot-and-mouth disease (FMD) is a highly contagious disease that spreads among buffalo populations, and a major concern of FMD is in how persistently infected hosts can cause potential obstacles in disease eradication in South African territories. We formulate a novel immuno-epidemiological model of FMD in African buffalo to investigate the role of carrier transmission on the disease dynamics. This model takes both the acute and carrier stages of the disease into consideration by incorporating two within-host models that capture the viral and immune dynamics within an FMD infected buffalo experiencing only acute or both phases and by coupling these models to a time-since-infection age structured epidemiological model.

In analyzing the within-host model of carrier-infected buffalo, we define an immunological basic reproduction number, ℛ_0_, that also serves as a threshold condition for the local stability of a pathogen-free equilibrium. Additionally, we present existence and stability conditions for two pathogen persisting equilibria. Furthermore, we analyze the immuno-epidemiological multi-scale model. We define a within-host viral-immune kinetics dependent basic reproduction number, ℛ_0_, and later show that it serves as a threshold condition for the local stability of the disease-free equilibrium and the existence of the endemic equilibrium. We also analytically show that the epidemiological model always displays forward bifurcation for specific between-host parameters, excluding the possibility of backward bifurcation.

Numerical results indicate that two mechanisms drive the persistence of FMD. One mechanism pertains to the role of FMDV persistence due to carrier-infected buffalo remaining in the population. The second mechanism is that FMDV operates as a childhood disease in the sense that carrier and acute infected populations of a specific strain remain in the population where they can introduce that strain to newly birthed calves. Such results align with Jolles et al. (2021). We apply sensitivity analysis to better understand the impact of within-host parameters on ℛ_0_. Sensitivity analysis suggests that the most efficient control measures are the ones that prioritize boosting of the adaptive immune response, and such measures should be applied even while individuals are carriers. Through our sensitivity analysis results, we also recognize that solutions to the within-host carrier model can display viral relapse for varying values of the parameters. Viral relapse could be one of the reasons why buffalo remain infected for longer periods of times.

Further analysis on foot-and-mouth disease can be useful in elucidating how fast transmitting and evolving diseases spread among a population and how to manage such outbreaks. A future research endeavor of ours is to validate the immunological carrier model and the multiscale model with accross scale data via practical identifiability of model parameters. We are also interested in formulating and analyzing nested immune-epidemiological coinfection models for FMD and in investigating the application of immunological control strategies by applying optimal control theory to the within-acute and within-carrier model, where we can also investigate the epidemiological impacts that such strategies have on curtailing the spread of the disease.

## 7 Data availability

Not applicable.

## 8 Acknowledgements

We would like to thank Anna Jolles (Oregon State University), Brianna Beechler (Oregon State University), Joshua Caleb Macdonald (Tel Aviv University), Simon Gubbins (The Pirbright Institute), Jan Medlock (Oregon State University), and Maia Martcheva (University of Florida) for conversations that were helpful to us while conducting this research.

## Appendix A Analyses of the within-carrier model

In analyzing the within-carrier system (2), we assume *a* = 0 to align with results in fitting the system. Thus, the equilibria of the within-carrier system are given by the solutions of

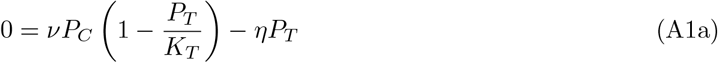

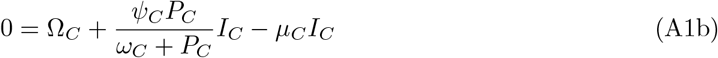

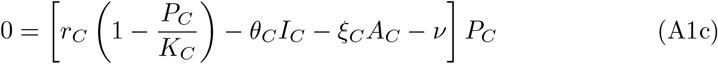

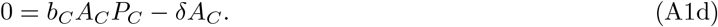

The generic Jacobian of the system is

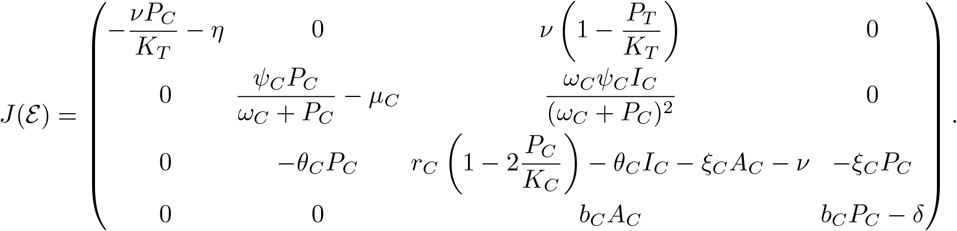

The computed Jacobian presented above will be used for proving Theorems 2.1-2.3, which will be presented in the following subsections.

### A.1 Stability analysis of the pathogen-free equilibrium

#### Theorem 2.1.

*The pathogen-free equilibrium*

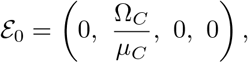

*which always exists, is locally asymptotically stable when*

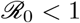

*and unstable when*

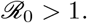

*Proof*. The Jacobian evaluated at ℰ_0_ is

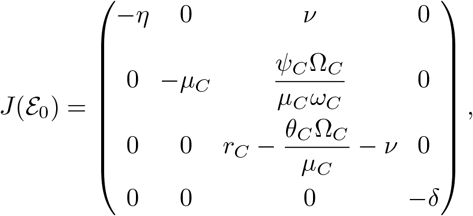

which has eigenvalues *λ*_1_ = *−η, λ*_2_ = *−μ*_*C*_, *λ*_3_ = *−δ*, and 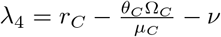. Since *λ*_1_, *λ*_2_, *λ*_3_ < 0, the stability of ℰ_0_ is determined by the sign of *λ*_4_.

We can rewrite *λ*_4_ as follows:

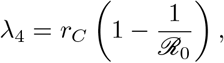

Where

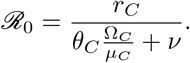

Therefore, ℰ_0_ is locally asymptotically stable when

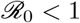

and unstable when

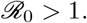

### A.2 Stability analysis of the pathogen-persisting boundary equilibrium

We next consider the possibility of a boundary pathogen-persisting equilibrium 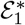. In this case, the adaptive immune response is absent, i.e., 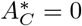. From (A1a), we have

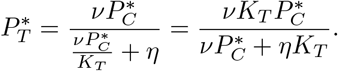

From (A1b), we have

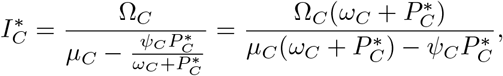

which exists when 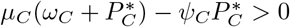. Substituting the above expression into (A1c), we obtain

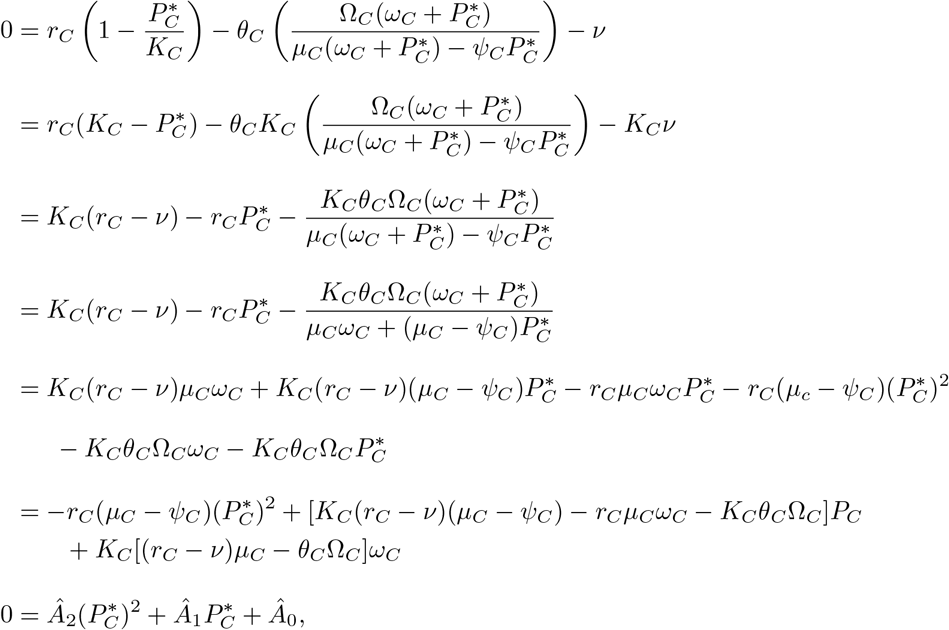

Where

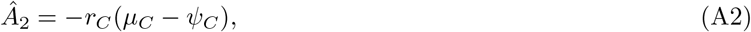

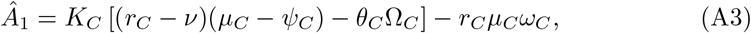

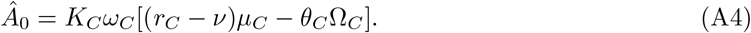

We denote *f* (*P*_*C*_) as being the following quadratic polynomial

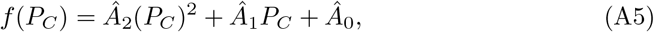

with coefficients being defined in equations (A2)-(A4). Before discussing the theorem of existence, uniqueness, and stability of a pathogen-persisting boundary equilibrium, we present the following lemma.

#### Lemma A.1.

*The following hold true:*

*Case 1: If* ℛ_0_ *<* 1 *and μ*_*C*_ *> ψ*_*C*_, *then Equation (A5) has no positive roots*.

*Case 2: If* ℛ_0_ *<* 1 *and μ*_*C*_ *< ψ*_*C*_, *then Equation (A5) has one positive root*.

*Case 3: If* ℛ_0_ *>* 1 *and μ*_*C*_ *> ψ*_*C*_, *then Equation (A5) has one positive root*.

*Case 4: If* ℛ_0_ *>* 1 *and μ*_*C*_ *< ψ*_*C*_, *then Equation (A5) has either two positive roots or no positive roots*.

*Proof*. Verification of each case follows directly from Descartes’ Rule of Signs and a couple of observations. First we can rewrite *Â*_1_ and *Â*_0_ as follows:

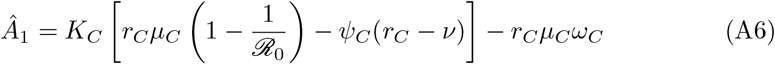

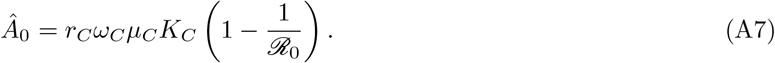

Additionally, we have that ℛ_0_ *>* 1 implies that *r*_*C*_ *> v*.

**Case 1**: Assume ℛ_0_ *<* 1 and *μ*_*C*_ *> ψ*_*C*_. Then, it follows that *Â*_2_ *<* 0 and *Â*_0_ *<* 0. For determining the sign of *Â*_1_ we consider two subcases.

**Subcase 1a)** Assume *r*_*C*_ *< v*. From Equation (A3) the term (*r*_*C*_ *− v*)(*μ*_*C*_ *− ψ*_*C*_) *<* 0. This allows us to deduce that *Â*_1_ *<* 0, and (by Descartes’ Rule of Signs) Equation (A5) has no positive roots.

**Subcase 1b)** Assume *r*_*C*_ *≥ v*. From Equation (A6) we have *r*_*C*_(1 *−* 1*/*ℛ_0_) *<* 0 since ℛ_0_ *<* 1 and the other two terms are clearly negative. We can deduce that *Â*_1_ *<* 0, so Equation (A5) has no positive roots.

**Case 2)** Assume ℛ_0_ *<* 1 and *μ*_*C*_ *< ψ*_*C*_. Then, *Â*_2_ *>* 0 and *Â*_0_ *<* 0. We can then deduce that Equation (A5) has exactly one positive root.

**Case 3)** Assume ℛ_0_ *>* 1 and *μ*_*C*_ *> ψ*_*C*_. It then follows that *Â*_2_ *<* 0 and *Â*_0_ *>* 0, which allows us to deduce that Equation (A5) has exactly 1 positive root.

**Case 4)** Assume ℛ_0_ *>* 1 and *μ*_*C*_ *< ψ*_*C*_. Then, it follows that *Â*_2_ *>* 0, *Â*_0_ *>* 0, and *Â*_1_ *<* 0. By Descartes’ Rule of Signs Equation (A5) either has two positive roots or no positive roots. □

#### Theorem 2.2.

*If one of the two following conditions below holds:*

1. ℛ_0_ *>* 1 *and μ*_*C*_ *> ψ*_*C*_.
2. ℛ_0_ *>* 1, *μ*_*C*_ *< ψ*_*C*_, *and the discriminant of the quadratic polynomial f* 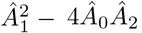, *is positive*,
3. *then there exists a unique pathogen-persisting boundary equilibrium* 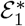. *Moreover, if* 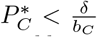, *then the pathogen persisting boundary equilibrium is locally asymptotically stable*.

*Proof*. Before discussing conditions 1 and 2, we need to verify that the parameters set to satisfy cases 1 and 2 from Lemma A.1 (the cases in which ℛ_0_ *<* 1) imply that the pathogen-persisting boundary equilibrium does not exist. In order for the pathogen-persisting boundary equilibrium to exist, we need 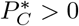 and 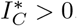. In Lemma A.1, we showed that if ℛ_0_ *<* 1, and *μ*_*C*_ *> ψ*_*C*_, there is no positive root 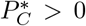, so no boundary equilibrium arises from this case. In Lemma A.1, we showed that if ℛ_0_ *<* 1 and *μ*_*C*_ *< ψ*_*C*_, then we have existence of exactly one positive root 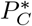. In order for the steady state value 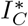 to also be positive, 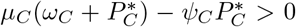 or equivalently 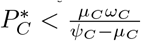. We can use properties of the graph of the quadratic polynomial *f* and the value of 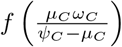 to show that this inequality does not hold when ℛ_0_ *<* 1 and *μ*_*C*_ *< ψ*_*C*_.

As mentioned in the proof of Lemma A.1, ℛ_0_ *<* 1 and *μ*_*C*_ *< ψ*_*C*_ implies that *Â*_2_ *>* 0 and *Â*_0_ *<* 0. This implies that the graph of *f* is a parabola that opens up with *y*-intercept below the *x*-axis. Thus showing that the positive root of *f* satisfies 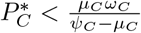 is equivalent to showing that 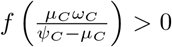. We evaluate 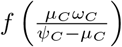 with (A7), (A6), and (A2) substituted in for *Â*_0_, *Â*_1_, and *Â*_2_ (respectively) and use algebra to simplify terms to get

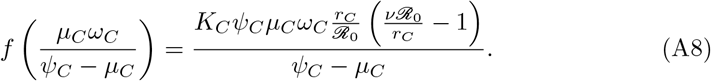

Note that the sign of the numerator depends on the sign of (*v*ℛ_0_*/r*_*C*_) *−* 1. We use (3) to obtain:

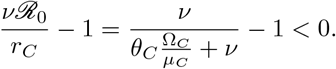

Thus, the numerator for 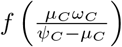 is always negative, and this holds true regardless of whether or not ℛ_0_ is greater than or less than one. Since 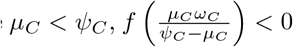. Therefore, the positive root corresponding to the case in which ℛ_0_ *<* 1 and *μ*_*C*_ *< ψ*_*C*_ does not satisfy 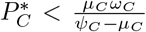 which allows us to deduce that no boundary equilibrium exists in this scenario.

In Lemma A.1, we showed that if ℛ_0_ *>* 1 and *μ*_*C*_ *> ψ*_*C*_ (this is Condition 1 from our theorem), then *f* has a unique positive root 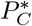 . In order for the steady state value 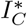 to also be positive in this case, 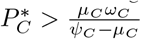, which is immediately satisfied since the right hand side of the inequality is negative.

In Lemma A.1, we showed that ℛ_0_ *>* 1 and *μ*_*C*_ *< ψ*_*C*_ implies that *f* has either two positive roots or no positive roots. In order to have positive roots, the discriminant of the quadratic polynomial *f*, that is 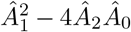, must be positive, which is why such an inequality is added to condition 2 of our theorem. Although we get the existence of two positive roots in this scenario, the smaller positive root, which we denote as 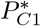, is the only root that yields a positive steady state value for 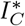. In order for 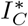 to be positive, we need it to be the case that 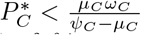. If the parameters are set to satisfy condition 2 of the theorem, the graph of *f* is a parabola that opens up with the *y*-intercept lying above the *x*-axis. We denote 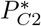 as being the larger positive root.

Based upon the characteristics of *f*, the graph of *f* (*x*) should lie below the *x*-axis for any 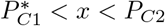 and the graph of *f* (*x*) is above the *x*-axis for any 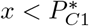 and any 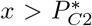. From Equation (A8) we have that 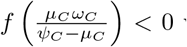 when *μ*_*C*_ *< ψ*_*C*_, which implies that 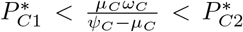 . Therefore, the steady state value of the innate immune response, 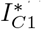, corresponding to the smallest positive root, 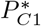, is positive while the computed value of the innate immune response, 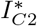, corresponding to the larger positive root, 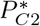, is negative. This tells us that for condition 2 we get existence of a unique pathogen persisting boundary equilibrium.

To analyze the stability of 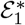 we compute the Jacobian

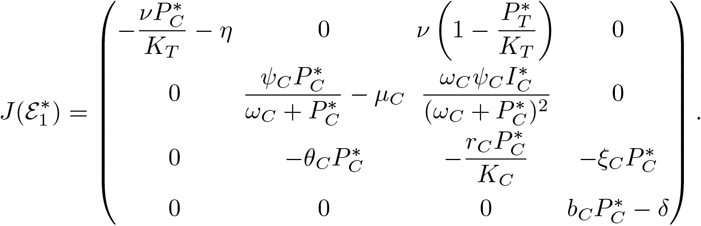

The matrix 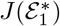 has two eigenvalues

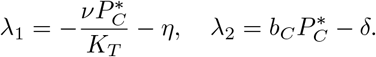

The sign of *λ*_1_ is negative and the sign of *λ*_2_ is negative if 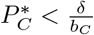 . To investigate the remaining eigenvalues, we first observe that the second diagonal entry of 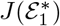 can be rewritten as 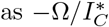, since 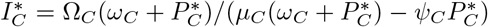.Then we consider the following submatrix

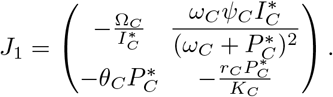

We can determine the signs of the remaining eigenvalues by analyzing the trace of *J*_1_ and the determinant of *J*_1_ . Observe 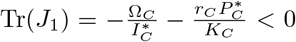 and det 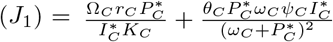, which implies that the sign of the remaining eigenvalues are negative. This completes our proof.

### A.3 Stability analysis of the pathogen-persisting interior equilibrium

We now consider the pathogen-persisting interior equilibrium

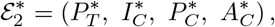

with components

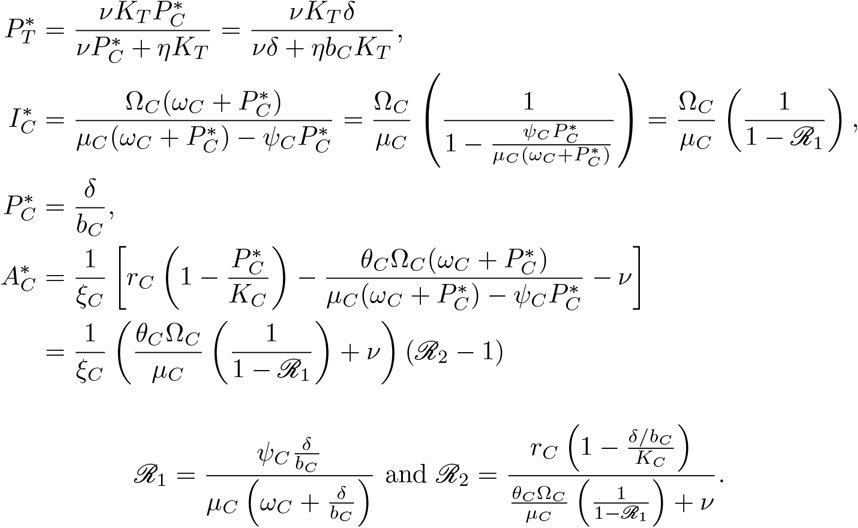

Where

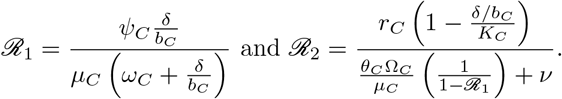

#### Theorem 2.3.

*The pathogen-persisting interior equilibrium*, 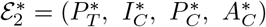, *exists and is locally asymptotically stable if*

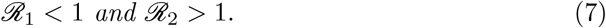

*Proof*. The condition *ℛ*_1_ *<* 1 ensures that 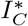 is positive, and the second condition *ℛ*_2_ *>* 1 in the theorem ensures that 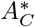is positive. We now linearize (2) about this equilibrium. First, note that as *P*_*C*_ *≠* 0, (A1c) implies

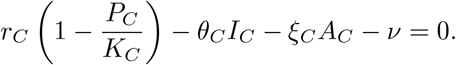

Thus, the Jacobian evaluated at 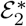is

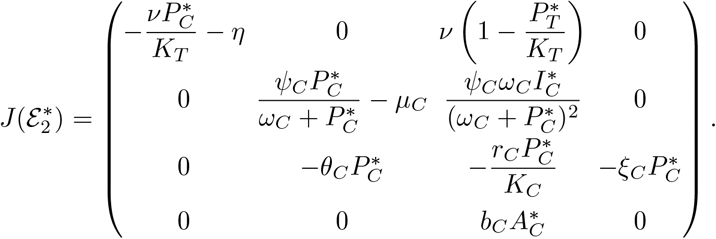

One of the eigenvalues of 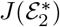is

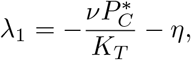

which is negative.

We use *ℛ*_1_ to rewrite the second diagonal entry of 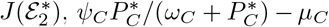, as being *−μ*_*C*_(1 *− ℛ*_1_). Consider the submatrix

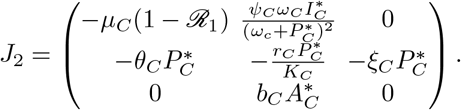

The remaining eigenvalues are the solutions of the following equation

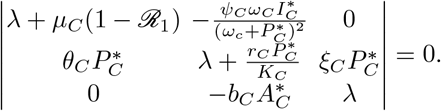

By expanding the determinant, we have

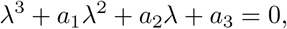

Where

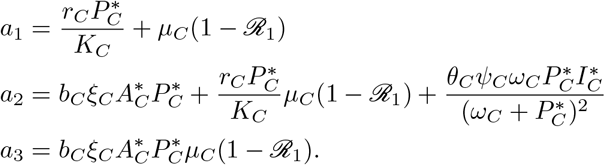

We now apply the Routh-Hurwitz criterion to show that the steady state is locally asymptotically stable. Coefficients *a*_1_, *a*_2_, and *a*_3_ are positive if *ℛ*_1_ *<* 1 and *ℛ*_2_ *>* 1 (the latter ensures that 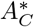 used in *a*_2_ and *a*_3_ is positive). Observe

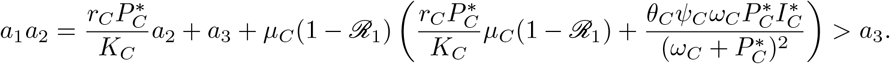

All of the conditions for the Routh-Hurwitz Criterion are satisfied which implies that the roots of the characteristic equation have negative real parts. This completes our proof.□

## Appendix B The numerical scheme

For the epidemiological model, we approximate solutions to the age-structured PDE model with the stored within-host calculations. Let 0 *≤ t ≤ T* be the time interval of interest and 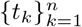 be a partition of the time interval with fixed mesh size Δ*t* = *T/n*. Let 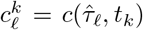 and 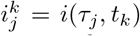. Let 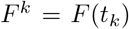 for all *F ∈ {M, S, C, I, R, N}*.The numerical scheme, a mixed implicit-explicit scheme, is as follows:

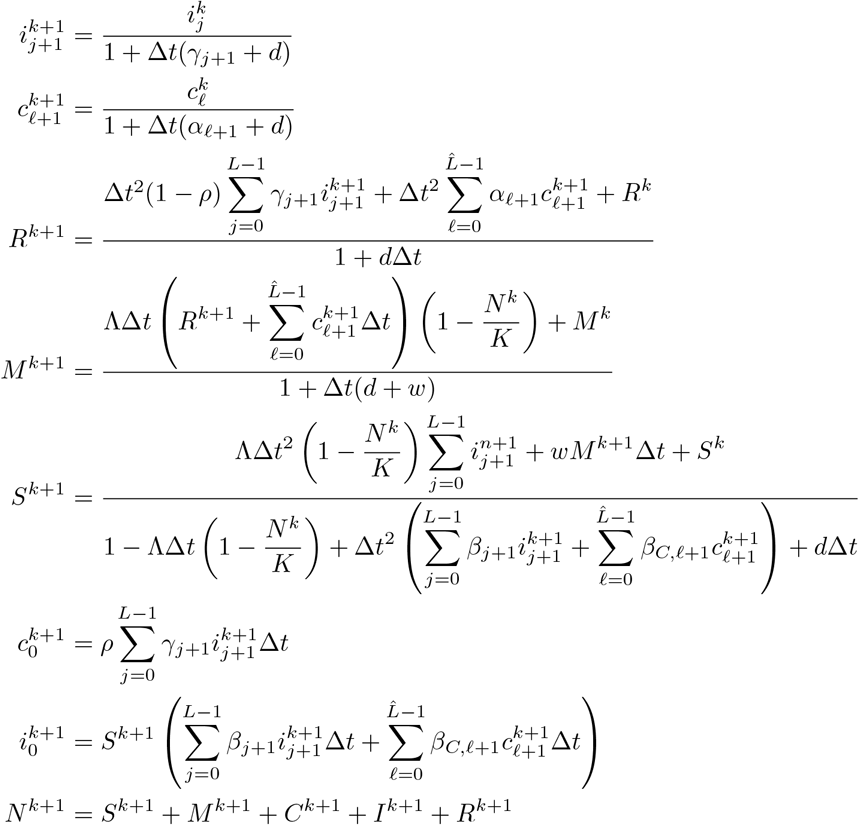

where *n, L*, and 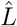 are chosen large enough to where 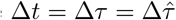.

## Appendix C Sensitivity analysis

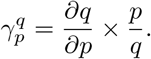

In our case, *q* is the basic reproduction number *ℛ*_0_ for the original system (8), *p* is any within-host viral and immune kinetics parameter during either the acute or carrier phase (*i*.*e. p* can be the viral growth rate parameter *r*_*A*_).

This approach can be implemented across the scales for different QOIs by defining linking functional forms. Through the linking functions appearing in the construction of *ℛ*_0_, *ℛ*_0_ depends implicitly on the within-host state variables. Let *p* be some parameter used in the within-host model for an infectious individual during the acute phase. The partial derivative of *ℛ*_0_ with respect to the within-host parameter *p* is the following:

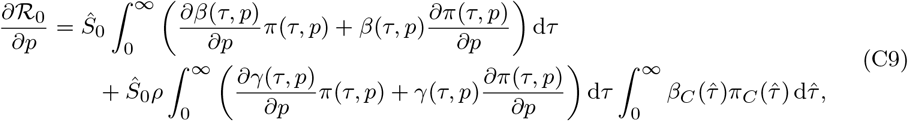

where

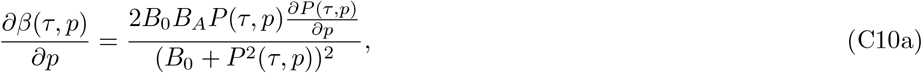

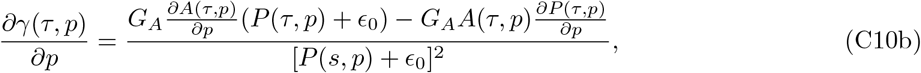

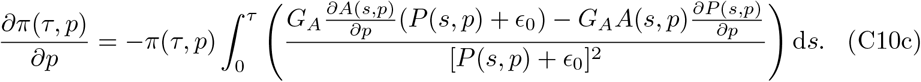

Let 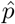 be some parameter used in the within-host model for an infected individual during the carrier phase. The partial derivative of *ℛ*_0_ with respect to the within-carrier parameter 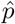 is as follows:

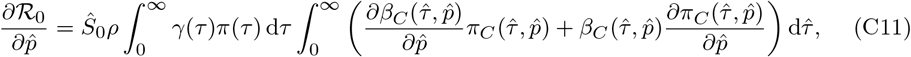

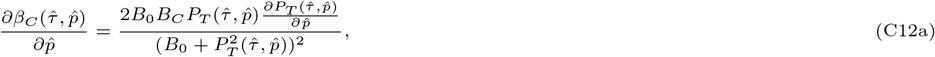

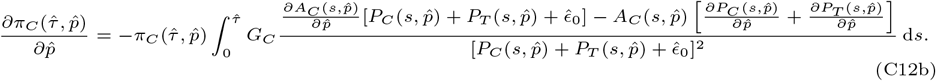

Observe that we need to compute 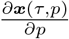 and 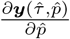 where ***x***(*τ, p*) = (*I*_*A*_(*τ, p*), *P* (*τ, p*), *A*(*τ, p*)) and 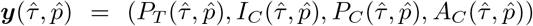. By Clairaut’s Theorem, we have that

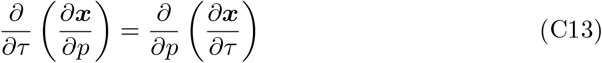

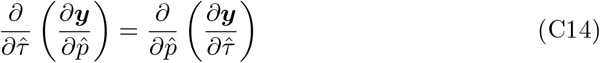

where *∂****x****/∂τ* is the right hand side of system (1) and *∂****y****/∂τ* is the right-hand side of system (2). Let ***f*** (***x***, *p*) = *∂****x****/∂τ*, and consider the following system for the derivatives *∂****x****/∂p*:

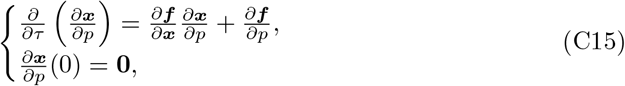

where *∂****f*** */∂****x*** is the Jacobian matrix of the right-hand side of the system (1). We find the solutions for ***x*** and the derivatives *∂****x****/∂p* at all times *τ*, by solving the extended system (C15)-(1) and we use these solutions to estimate the integrands used in (C9) and (C12b). To numerically solve the extended system (C15)-(1), we use ode45 with mesh size set to 0.01. Additionally, we have the relative and absolute error tolerances for the ODE solver set to 10^*−*10^ and 10^*−*11^, respectively. For approximating the integrals used in (C9) and (C12b), we use the trapezoidal rule.

We incorporate the same techniques for finding 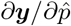. Let 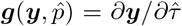and consider the following system:

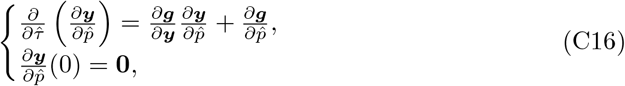

*∂****g****/∂****y*** being the Jacobian matrix of the right-hand side of the system (2).

Here are a few biological assumptions implemented in the numerical computation of SA:

- *For the computations in (C9) and (C12b), the approximated limits of integration that are used for the integrals associated with the acute phase is* 0 *and* 30. *Observe in Figure 2, the pathogen load clears well before day 30 and the adaptive immune response is already stabilizing to a fixed value by day 30. For the integrals associated with the carrier phase, we use 30 and 750 as our lower and upper bounds of integrals*.
- *The initial conditions used for the within-carrier model are the same values as the ones that are used for acute phase, which leads to a re-simulation of an acute phase. In our previous simulations of the multiscale model we ignore the first τ*_*\**_ *days of the carrier phase (where τ*_*\**_ *is the time for which the pathogen load in the bloodstream P*_*C*_(*τ*_*\**_) *<* 10^*−*4^*); however, for the sensitivity analysis experiments, we ignore the first thirty days of the carrier phase since the 28-30 day mark (post infection) is often considered as being the conventional start time of the carrier phase (Stenfeldt C, 2020). Additionally, we observe that in varying some of the within-carrier parameter values (in particular parameters v, K*_*C*_, *r*_*C*_, *δ), we have earlier instances of viral relapse during the carrier phase (in which case τ*_*\**_ *cannot be numerically obtained). We provide plots of these instances in Figure 10. Viral relapse is also observed in generated experimental data, used for the study Macdonald et al. (2022). The computed R*_0_ *value at the baseline parameters is 21*.*8516*.

## Appendix D Additional figures

